# Accurate ab *initio* gene prediction in eukaryotes with Tiberius in multiple clades

**DOI:** 10.64898/2026.04.24.720536

**Authors:** Lars Gabriel, Tomáš Brůna, Asees Kaur, Anish Krishnan, Felix Ortmann, Asaf Salamov, Samuel Talbot, Felix Becker, Richard Krieg, Christopher W. Wheat, Igor V. Grigoriev, Mario Stanke, Katharina J. Hoff

**Author notes:** These authors contributed equally as senior authors (Mario Stanke, Katharina J. Hoff).

## Abstract

**Background:** Eukaryotic genome annotation is currently bottlenecked by limitations in the generality, scalability or accuracy of computational methods and fewer than 20% of the genomes available at NCBI Datasets have associated gene annotations. Evidence-based pipelines such as BRAKER3 achieve high accuracy but require substantial extrinsic evidence and compute. Deep learning approaches have recently achieved large improvements in *ab initio* gene prediction accuracy. The end-to-end deep learning gene predictor Tiberius approaches the accuracy of evidence-based annotation on mammalian genomes without using any extrinsic evidence, but its published models were trained on mammals only, limiting its applicability across eukaryotes.

**Results:** We extend Tiberius beyond mammals by training lineage-specific models for Mesangiospermae, Fungi, Vertebrata, Insecta, Chlorophyta and Bacillariopliyta, covering clades that account for 92% of currently available eukaryotic assemblies. Across a benchmark of 33 species, Tiberius achieved higher exon-, transcript- and gene-level accuracy than the other evaluated *ab initio* methods, Helixer and ANNEVO, improving gene-level F1 score by 12-37 percentage points over Helixer and by 10-22 percentage points over ANNEVO, while also having the fastest runtimes overall. Compared with BRAKER3, which incorporates RNA-Seq and protein evidence, Tiberius approaches state-of-the-art accuracy in Mesangiospermae, Fungi, Bacillariophyta and Chlorophyta, while being on average 80 times faster when using a GPU. A reimplementation of the Tiberius backend reduced runtime by 31% compared to the previous implementation. Tiberius and its models are also available through a web server, which allows annotation of submitted assemblies without local resources. Furthermore, the Vertebrata model has been applied to annotate 2,948 vertebrate assemblies totalling nearly 6 trillion base pairs.

**Conclusions:** Tiberius is transferable well beyond Mammalia and reaches accuracy close to evidence-based annotation in several clades at a small fraction of the compute cost. This makes it a practical choice for highly accurate large-scale genome annotation, particularly when extrinsic evidence is unavailable or annotation throughput is limiting.

**Availability and implementation:** Code: https://github.com/Gaius-Augustus/Tiberius, web server: https://bioinf.uni-greifswald.de/tiberius.

## 1 Introduction

Accurate Identification of protein-coding genes is a critical step in genome annotation, yet fewer than 20% of the currently available genomes at NCBI Datasets [1] have associated gene annotations. This problem may amplify quickly as the scale of genome sequencing continues to expand through initiatives such as the Earth BioGenome Project [2], Moreover, since non-uniform genome annotations in comparative genomic analyses can dramatically inflate estimates of lineage-specific genes [3], there is a growing need for tools that can rapidly generate uniform genome annotations for large genomic datasets. Improving accuracy, ease of application and compute efficiency is therefore essential to homogeneously annotate the growing wealth of available genomes.

While evidence-based annotation methods that integrate RNA-Seq data and protein homology, such as BRAKER3 [4], achieve high accuracy, they require substantial extrinsic evidence that may not be available for many newly sequenced organisms. In addition, BRAKER3’s computational requirements can be prohibitive when annotating large numbers of genomes; this is exacerbated by the need for generating species-specific repeat libraries for rigorous repeat masking prior to gene annotation.

Until recently, *ab initio* gene prediction methods, which identify genes solely from the genome sequence, have severely lagged behind evidence-based approaches in terms of accuracy. Additionally, many *ab initio* gene predictors, such as AUGUSTUS [5], require species-specific training, which is time-consuming and requires extrinsic evidence such as RNA-Seq data or homologous proteins to achieve decent accuracy.

Helixer [6], introduced in 2023, showed that combining deep learning layers and hidden Markov model (HMM) postprocessing could improve over the “shallow” HMM gene-finder AUGUSTUS. Building on the tindings of the Helixer project, we recently introduced Tiberius [7], an end-to-end deep learning *ab initio* gene predictor that achieves high accuracy for mammalian genomes, approaching the accuracy of BRAKER3 despite using no extrinsic evidence. More recently, ANNEVO has been introduced as another deep learning-based *ab initio* gene predictor that outcompetes Helixer in terms of accuracy [8],

The original Tiberius was trained only on mammals, limiting its applicability across eukaryotes. Here, we extend Tiberius to six additional clades: Mesangiospermae, Fungi, Vertebrata, Insecta, Chlorophyta and Bacillariophyta, which cover 92% of currently available eukaryotic assemblies at NCBI (Figure 1). We show that these models outperform Helixer and ANNEVO across almost all benchmarks while approaching BRAKER3 accuracy in several lineages.

**Figure 1:**
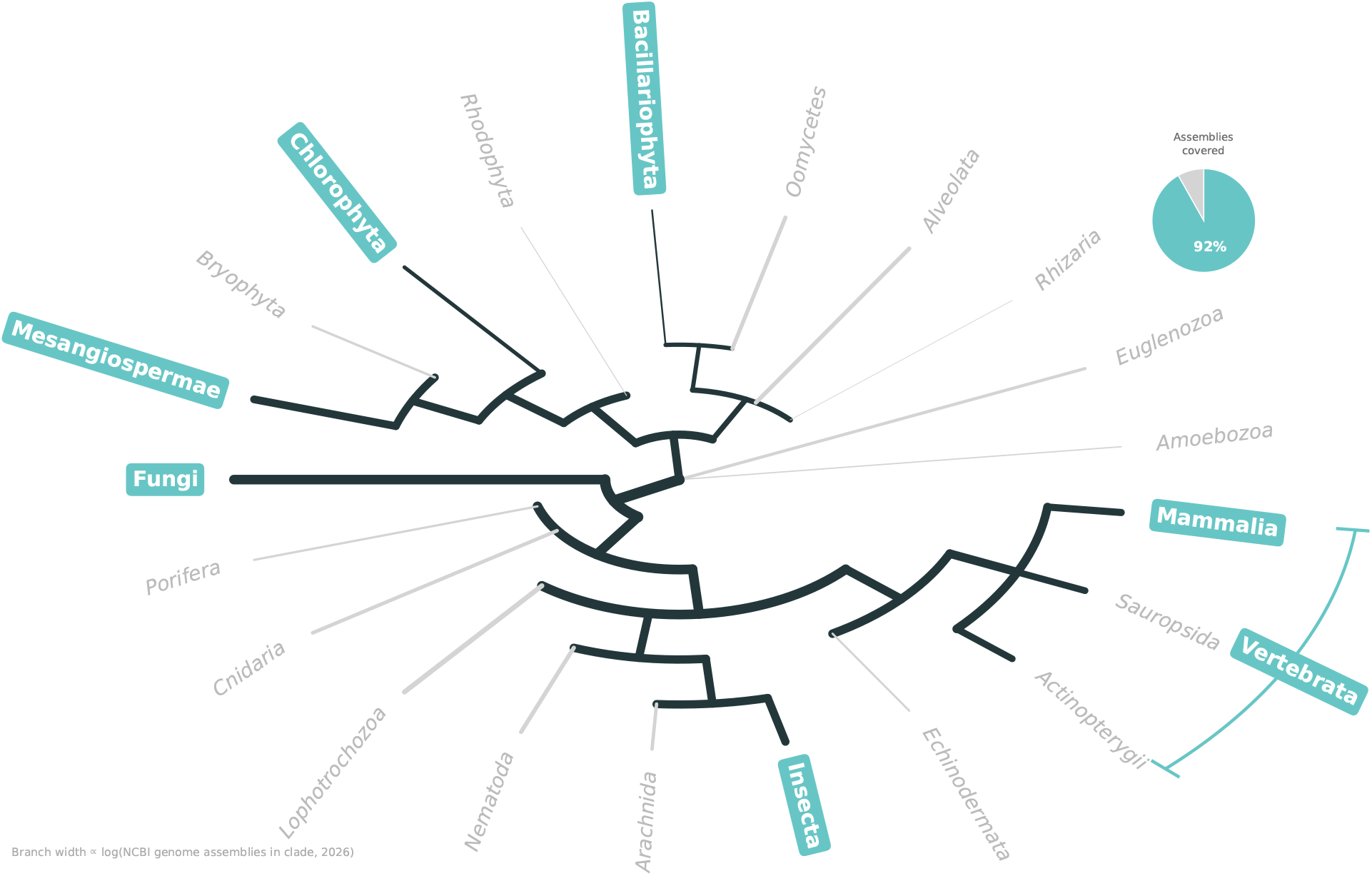
Taxonomic clade space covered by the Tiberius models. Schematic phylogeny of major eukaryotic groups, the branch width is proportional to the logarithm of the number of genome assemblies available for the respective clade at NCBI. Clades with a Tiberius model are highlighted in teal. Groups in grey are not covered by a cladc-spccific Tiberius model. The inset shows the proportion of eukaryotic assemblies at NCBI belonging to a covered clade, weighted by assembly count rather than by number of species (55,786 of 60,750 assemblies, 91.8%). Data retrieved from NCBI Datasets [1] on 2026-03-25.

## 2 Materials and methods

### 2.1 Model training

Clade-specific Tiberius models were trained for Mesangiospermae, Fungi, Vertebrata, Insecta, Chlorophyta and Bacillariophyta. All models were trained on unmasked genomes (training, validation, test data see Supplementary Materials: Tables S1–S9) following the procedure described in the original Tiberius publication (see Supplementary Materials). Details on hardware and model training are provided in Supplementary Materials: Tables S29–S30.

The overall architecture remained unchanged, except for adjusted layer widths in Insecta and Vertebrata (Supplementary Materials: Table S30).

### 2.2 Implementation improvements

The Python backend of Tiberius was mostly reimplemented from scratch and modularized into two separate packages. The first package, *bricks2marble*, is a fast and versatile framework for genomic data handling and annotation logic that is easily extendable. It covers sequence encoding, windowing and the reading and writing of annotation bles independently of a particular model architecture. The second package, *hidten*, provides the differentiable HMM layer as a standalone component, independently of its State space. Both packages are usable independently of Tiberius for future genome processing tools. In addition, the new implementation is compatible with more recent TensorFlow versions (>2.13), making it easier to install and maintain.

### 2.3 Benchmark comparison

We benchmarked Tiberius against the deep learning *ab initio* gene prediction tools, Helixer and ANNEVO. For each tool, clade-appropriate pretrained models provided by the respective authors were used where available (Supplementary Materials: Table S28). In addition, we compared performance with that of BRAKER3 as a representative state-of-the-art general-purpose gene prediction pipeline. In contrast to the *ab initio* tools, BRAKER3 requires transcriptomic and protein evidence. RNA-Seq input was generated using VARUS [9], As protein evidence, we used the combined proteomes of all Tiberius training species within the respective target clade. Across all clades, 37 test species were used to evaluate Tiberius. Of these, 33 were benchmarked against all four tools. The remaining two clades were cxcludcd from the overall comparison because pre-trained models were unavailable for some of the tools: there was no ANNEVO model for Chlorophyta or Bacillariophyta and no Helixer model for Bacillariophyta. Reference annotations of the test species were obtained from NCBI RefSeq, Ensembl, GenBank, Phytozome or the respective genome projects (Supplementary Materials: Tables S1–S9). None was generated by our group, but two were generated by other groups with BRAKER2: the *E. debaryana* annotation [10] and the *T. pseudonana* annotation, which was subsequently corrected with PASA [11], Neither species is part of the 33-species comparison of all four tools.

### 2.4 Evaluation metrics

Prediction accuracy was evaluated using standard metrics for gene prediction. Predicted gene structures were compared to reference annotations at the coding sequence (CDS) exon and gene level, with untranslated regions (UTRs) excluded from the evaluation. A predicted instance was considered a true positive only if its structure exactly matched a reference instance, except that the outer ends of the coding region (start and stop codon ends) were allowed to deviate by up to 3bp to accommodate differing conventions for including the stop codon in CDS features (`gffcompare -strict-match -e 3`; Supplementary Materials). Throughout this manuscript, “exact” refers to this criterion. At the gene level, a locus was counted as correctly predicted if at least one transcript matched the reference annotation exactly. Sensitivity, precision and F1 score were calculated from the resulting counts using `gffcompare` v0.12.10 [12],

## 3 Results

### 3.1 Accuracy comparison with other *ab initio* methods

On all five clades (33 test species) for which all three tools were trained, Tiberius achieved higher exon-, gene- and transcript-level accuracy than Helixer and ANNEVO (Figure 2). At gene level, Tiberius improved F1 scores over Helixer on average by 12–37 percentage points per clade, with the largest differences in Mammalia and non-mammalian Vertebrata (Supplementary Materials: Tables S18–S25). Compared with ANNEVO, Tiberius achieved a 10–22 percentage points higher gene-level F1 score, with higher differences in sensitivity than precision. Tiberius’ exon-level F1 score was also consistently higher, by about 5 percentage points on average. Tiberius achieved a higher F1 score than both Helixer and ANNEVO on each of the 33 test genomes at exon, transcript and gene level. Pairing per genome (n =33 genomes, one F1 value per tool per genome), two-sided Wilcoxon signed-rank tests confirmed that these per-genome differences are significant at all three levels for both comparators (Supplementary Materials: Table S31). Across the two Chlorophyta test species, Helixer achieved the higher gene-level F1 score on *B. prasinos* (55.8% vs. 50.1%), whereas Tiberius performed better on *E. debaryana* (39.3% vs. 30.3%). ANNEVO does not provide a pretrained model for Chlorophyta. Of the 33 test species, Helixer included 17 in its training or validation data, whereas ANNEVO included two and Tiberius included none (Supplementary Materials: Table S17). No test species shares a species with any Tiberius training species. Within the 33-species benchmark, only *D. melanogaster* shares a genus with a training species (28 Mya), all other test species are separated from the closest training species at family level or above, with divergence times of 14-281 Mya according to TimeTree [13] (Supplementary Materials: Table S32; no TimeTree estimate is available for three of these species). Proximity to the training data did not systematically predict accuracy. *U. brizantha* (14 Mya from the closest training species) was among the weakest Mesangiospermae results, whereas *E. californica* (108 Mya) was among the strongest. Tiberius offers the first deep learning gene prediction model for the protists Bacillariophyta. Therefore, on this clade, only comparisons to BRAKER3 were conducted.

**Figure 2:**
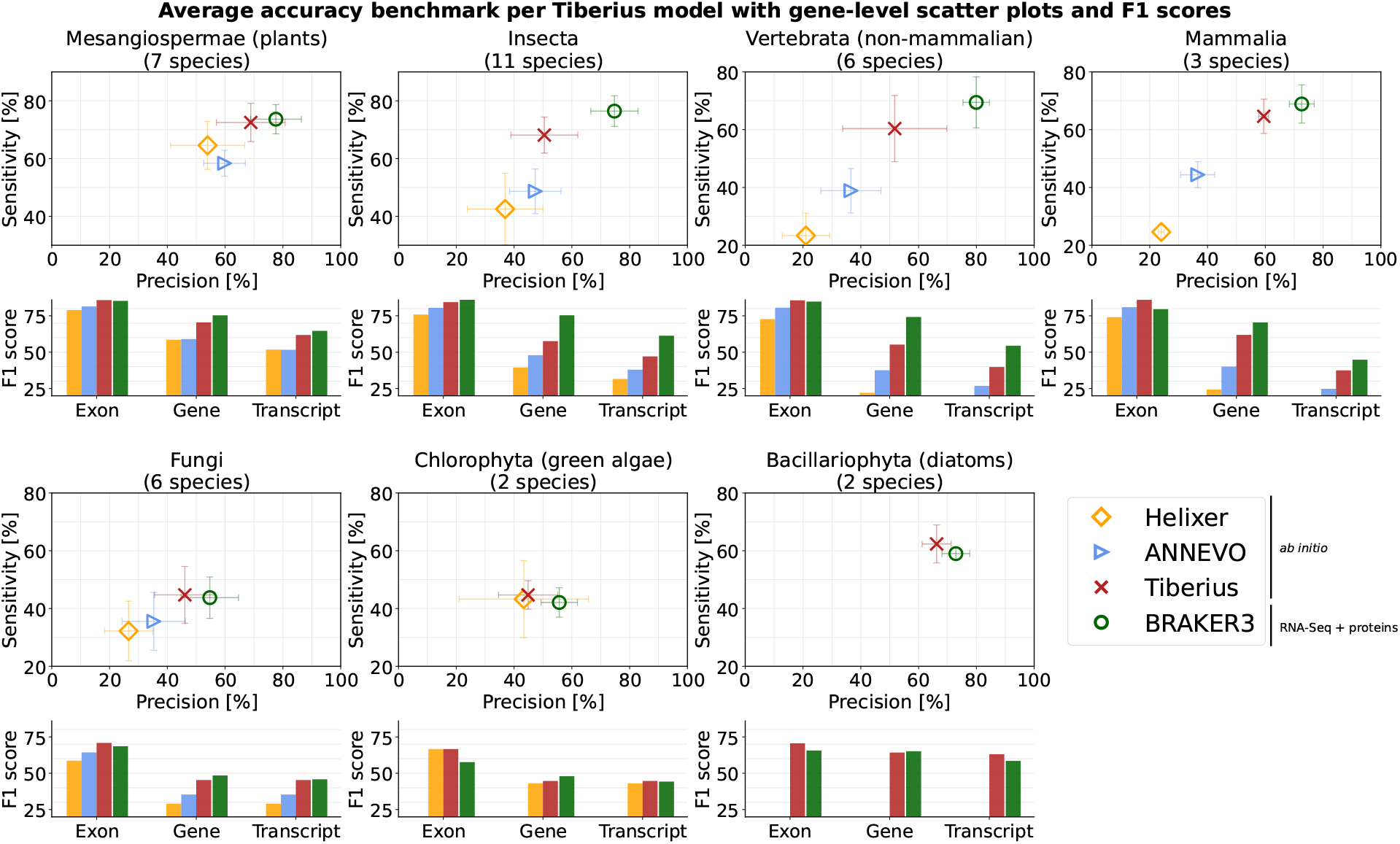
Average accuracy across test species for each Tiberius model, compared with *ab initio* predictions from Helixer and ANNEVO and evidence-supported predictions from BRAKER3. Gene-level sensitivity and precision are shown as scatter plots; CDS-exon-, transcript- and gene-level F1 scores are shown as bar plots. Markers show the unweighted mean across the test species of a clade (each species weighted equally, not pooled over genes); error bars show the standard deviation across these species (mean ± SD). F1 scores are computed from the mean sensitivity and precision. The number of test species per clade is given in the panel titles; per-species values are listed in Supplementary Materials: Tables S18–S25.

### 3.2 Accuracy comparison with BRAKER3

BRAKER3, which additionally uses RNA-Seq and protein evidence, achieved higher overall gene- and transcript-level accuracy than Tiberius, with overall F1 score differences of about 12 and nine percentage points, respectively. The largest gap was in non-mammalian Vertebrata, where BRAKER3 exceeded Tiberius by about 19 percentage points at gene level and 15 at transcript level F1 scores. In Mesangiospermae, Fungi, Bacillariophyta and Chlorophyta, transcript-level accuracy was similar and in Bacillariophyta, Tiberius exceeded BRAKER3 by fivc percentage points in F1 score. For these four groups, gene-level F1 scores differed by less than three percentage points. At exon level, both methods performed similarly, with Tiberius having on average an ≤ 1 percentage point higher F1 score.

### 3.3 Characterization of prediction errors

Aggregated F1 scores summarize accuracy but provide little information about the error patterns each tool makes, which may be important for downstream applications. We therefore characterized where the accuracy differences arise and how they affect predicted genes (Supplementary Materials: Figures S1–S5).

Stratifying gene-level accuracy by CDS length (Supplementary Materials: Figure S1) and CDS-exon count (Supplementary Materials: Figure S2) shows that most differences occur in long and multi-exon genes. Helixer and ANNEVO show the largest declines in gene-level sensitivity and precision for genes with a CDS length above 2 kb and generally with increasing CDS exon count, whereas Tiberius and BRAKER3 retain substantially higher accuracy in these categories. In Mammalia and non-mammalian Vertebrata, Tiberius and BRAKER3 perform similarly on shorter and less exon-rich genes, while BRAKER3 has a higher precision for the longest genes and those with many CDS exons. Very short genes (CDS ≤ 300 bp) are difficult for all four tools, which show low sensitivity and, owing to overprediction, low precision in this length range.

A notable exception to Tiberius’ overall higher accuracy compared to Helixer and ANNEVO is the prediction of single-exon genes in Insecta. While Tiberius achieves the highest sensitivity for these, its precision is markedly lower than that of all other tools (≈18% vs. ≈74% for ANNEVO), indicating substantial overprediction of single-exon genes for insects. BRAKER3 shows a smaller decrease in precision for this category. Notably, the Tiberius training data for Insecta were dominated by BRAKER annotations (Supplementary Materials: Table S7), which may have contributed to this pattern.

At the level of individual coding-exon boundaries, Tiberius matched the reference more often than Helixer and ANNEVO for the large majority of species at donor and acceptor splice sites and approached BRAKER3 despite using no extrinsic evidence (Supplementary Materials: Figure S3). Start- and stop-codon precision was more variable and Insecta was the one clade in which Tiberius had lower accuracy than the other tools on these two boundary types while being more accurate than them for predicting splice sites. This pattern is expected when single-exon genes are overpredicted, since a spurious single-exon gene contributes an incorrect start and stop codon but no splice sites.

We quantified fused reference genes (one predicted locus spanning more than one reference locus) and split reference genes (one reference locus covered by several predicted loci) (Supplementary Materials: Figure S4). Fusions were rare for all tools: with the exception of *P. striiformis*, fewer than *3%* of reference loci were fused in any genome. Among the *ab initio* tools, Tiberius had the lowest average fusion rate in every clade. Splits were more frequent. Tiberius split fewer reference genes than Helixer in every clade, but slightly more than ANNEVO overall (3.7% vs. 3.1%) and more than BRAKER3.

For every predicted gene with no reference overlap, we searched the translated sequence against UniProt SwissProt [14] as an indicator of whether such predictions resemble known proteins (Supplementary Materials: Figure S5). Tiberius produced by far the largest absolute number of novel loci with SwissProt homology. We define the hit rate as the percentage of a tool’s novel loci whose protein has a SwissProt hit at e-value ≤ 10^−10^ (DIAMOND blastp [15]). Tiberius’s hit rate was 30.6%, exceeding Helixer (26.0%), essentially tied with BRAKER3 (30.5%) and below ANNEVO (37.4%), which achieves a higher hit rate on a much smaller pool of novel predictions. We regard SwissProt homology only as a mild indicator. It shows that a prediction resembles a known protein, not that it is a functional gene and it does not distinguish genuine unannotated genes from pseudogenes and retrocopies. Hit rates are also not comparable across clades, since SwissProt underrepresents fungal and protist proteins.

Two test species stand out as difficult across these analyses. *Pristiophorus japonicus*, a Japanese sawshark, has by far the highest split rate in the benchmark (Supplementary Materials: Figure S4). It is also the largest test genome (5.9 Gb), the vertebrate test species most distant from any training species (225 Mya, Supplementary Materials: Table S32) and is as a shark genome exceptionally repeat-rich. Since Tiberius was trained on unmasked sequences, both phylogenetic distance and repeat content likely contribute to its low accuracy on this genome. *Puccinia striiformis*, the largest and most repeat-rich genome of the fungal test set, is the fusion-rate outlier (7.8% of reference genes fused).

### 3.4 Runtime comparison

A direct runtime comparison between BRAKER3 and the deep learning methods is limited because only the latter used a GPU. All tools used 72 threads of an AMD EPYC 7773X 64-Core Processor and the deep learning methods additionally used an NVIDIA A100-SXM4-80GB GPU. Across the 33 species benchmarked with all four tools, average runtimes were 26 min for Tiberius, 30 min for ANNEVO, 178 min for Helixer, and 2170 min for BRAKER3 (Table 1, Supplementary Materials: Tables S10–S16). BRAKER3 was particularly slow on large genomes, in non-mammalian Vertebrata it required on average 3035 min per genome, whereas Tiberius and ANNEVO remained below 60 min. Among the *ab initio* methods, Tiberius and ANNEVO were the fastest and about six times faster than Helixer on average.

**Table 1:** Average runtime per clade and overall mean across all species. The number of species per average is indicated as *n*. All tools used 72 threads of an AMD EPYC 7773X 64-Core Processor, the deep learning methods (Helixer, ANNEVO, Tiberius) additionally used an NVIDIA A100-SXM4-80GB GPU, whereas BRAKER3 ran on CPU only. Dashes indicate that no pretrained model was available for the respective clade. The overall mean covers the 33 species benchmarked with all four tools and therefore excludes Chlorophyta and Bacillariophyta.

| Clade | $n$ | GPU | | | CPU |
| --- | --- | --- | --- | --- | --- |
|  |  | Helixer | ANNEVO | Tiberius | BRAKER3 |
| Mesangiospermae | 7 | 1:35 | 0:22 | **0:17** | 20:26 |
| Insecta | 11 | 1:14 | **0:11** | 0:12 | 45:42 |
| Vertebrata | 6 | 6:27 | 1:10 | **0:58** | 50:35 |
| Mammalia | 3 | 11:00 | 1:32 | **1:21** | 77:08 |
| Fungi | 6 | 0:08 | **0:02** | **0:02** | 1:41 |
| Chlorophyta | 2 | 0:14 | – | **0:03** | 17:29 |
| Bacillariophyta | 2 | – | – | **0:01** | 19:13 |
| Overall | 33 | 2:57 | 0:30 | **0:26** | 36:05 |

### 3.5 Effect of implementation changes

The reimplementation reduced the average runtime per genome from 35 min to 24 min across all 37 test species, a 31% reduction (Supplementary Materials: Table S27), at matching prediction accuracy (Supplementary Materials: Table S26).

### 3.6 Large-scale vertebrate annotation

Tiberius has also been applied beyond the benchmark setting to annotate 2,948 vertebrate assemblies totaling nearly 6 trillion base pairs (5,987,909,637,336 bp) that are available online [16] (362 of primates, 1120 of other mammals, 497 of birds, 650 of fishes, 319 of other vertebrates). For 2,100 of these vertebrate assemblies the Tiberius annotations can be browsed in the UCSC Genome Browser [17, 18],

### 3.7 Web server

To make the clade-specific models available without local resources, we provide a web server [19], Users submit a genome assembly in FASTA format and select one of the available Tiberius models. The web server accepts gzipped assemblies of up to 500 Mb. Predictions are computed on either a GPU node (NVIDIA A100-SXM4-80GB GPU) or a CPU node (24 CPU cores and 63 GB of RAM) and returned as gene structures in GFF3 and GTF format. FASTA bles with predicted coding sequences and amino acid sequences are also provided. Results are retained for 45 days. The usage of the webserver does not require a registration. An email address can optionally be provided to receive a notification when the prediction is complete.

## 4 Discussion

### 4.1 Comparison with other *ab initio* methods

Our results show that Tiberius is transferable beyond Mammalia, and its models now cover clades that account for 92% of currently available eukaryotic assemblies (Figure 1). This percentage reflects the concentration of sequencing efforts in a few clades and is a measure of assembly coverage, not of species diversity. Coverage by a model also does not imply uniform accuracy: within a clade, accuracy varies with phylogenetic distance from the training data, repeat content and gene architecture (Section 3.3). Our benchmark also does not specifically test genomic features that are known to challenge annotation pipelines, such as trans-splicing, non-canonical splice sites, very short exons separated by long introns, very short introns, or combinations of these. Accuracy on genomes in which such features are frequent therefore remains to be established. Across this phylogenetic range, Tiberius was consistently more accurate than the other evaluated *ab initio* methods. Interpreting these comparisons is complicated by differences in training-data composition. Helixer, and to a lesser extent ANNEVO, included test species we selected for benchmarking in their training sets (Supplementary Materials: Table S17), giving them an advantage on these species [20].

For most clades, training data were assembled in collaboration with clade experts. The Fungi models are the exception, here both Tiberius and ANNEVO were trained on the same publicly available NCBI RefSeq annotations. This makes Fungi the most direct methodological comparison between the two tools. Even in this setting Tiberius achieved higher accuracy than ANNEVO across all metrics (Supplementary Materials: Table S22).

### 4.2 Model training

The variation in accuracy between clades suggests that reference annotation availability and quality remain major limiting factors. This is particularly evident for Chlorophyta, where training data are sparse and annotations are more heterogeneous than in better curated clades. For Insecta and Bacillariophyta, we supplemented publicly available annotations with BRAKER annotations generated by our group to improve clade coverage [21, 22], In Fungi, lower accuracy likely reflects broader phylogenetic diversity and heterogeneous annotations rather than insufficient training data. Further improvements will therefore require more consistently curated reference annotations across underrepresented clades.

### 4.3 Accuracy metrics

We evaluated predictions using exact (up to a 3-bp tolerance at the outer CDS ends, see Methods) CDS-exon-, transcript- and gene-level sensitivity, precision and F1 score, excluding UTRs because Tiberius does not predict them. These structure-level metrics have been traditionally used for benchmarking and remain the standard for gene prediction because they require correct reconstruction of exon boundaries and complete gene models [23, 24], Other measures, such as BUSCO completeness [25], nucleotide-level accuracy and protein-level similarity remain useful, particularly as a quality control measure for newly annotated genomes, but are not equivalent to exact structure-level assessment [7], These measures can miss biologically relevant errors in exon chaining and gene structure, which can affect downstream analyses [26, 27],

### 4.4 Comparison with BRAKER3

The comparison with BRAKER3 indicates that Tiberius’ main limitation is not CDS-exon-boundary detection, but the reconstruction of complete gene structures in complex genomes. This is supported by near parity of both methods at exon level, while a substantial gap remains at gene level in Vertebrata.

A limitation of the deep-learning *ab initio* methods evaluated here is that they predict only a single isoform per locus. This constrains transcript-level accuracy and gives BRAKER3 a systematic advantage at gene level, since a locus is counted as correct when at least one predicted isoform matches the reference. Extending deep-learning gene-finders to support multi-isoform prediction would improve biological realism. Another natural route to narrow the gap is the integration of extrinsic evidence, although this can come at substantial computational cost, as BRAKER3 required over 80 times more compute time than Tiberius.

## 5 Conclusion

We extend available Tiberius models from Mammalia to six additional clades which together account for 92% of currently available eukaryotic assemblies at NCBI. Tiberius consistently outperforms the other evaluated deep learning *ab initio* methods and, in several clades, approaches the accuracy of BRAKER3 while requiring far less runtime. These results make Tiberius a practical choice for highly accurate large-scale genome annotation, particularly when extrinsic evidence is unavailable or annotation throughput is limiting.

## 8 Data availability

Tiberius is open source and available at https://github.com/Gaius-Augustus/Tiberius under the MIT licence, version v2.0.0 was used for all analyses presented here and is archived at https://github.com/Gaius-Augustus/Tiberius/releases/tag/v2.0.0. The clade-specffic pretrained models are available from https://bioinf.uni-greifswald.de/bioinf/tiberius/models/angiosperms.tar.gz, https://bioinf.uni-greifswald.de/bioinf/tiberius/models/chlorophyta.tar.gz, https://bioinf.uni-greifswald.de/bioinf/tiberius/models/diatoms_unmasked.tar.gz, https://bioinf.uni-greifswald.de/bioinf/tiberius/models/fungi.tar.gz, https://bioinf.uni-greifswald.de/bioinf/tiberius/models/insecta_unmasked_v2.tar.gz, https://bioinf.uni-greifswald.de/bioinf/tiberius/models/tiberius_nosm_weights_v2.tar.gz and https://bioinf.uni-greifswald.de/bioinf/tiberius/models/vertebrates_weights.tar.gz. Tiberius and its models can be accessed also through a web server at https://bioinf.uni-greifswald.de/tiberius, which requires no registration.

The genome assemblies and reference annotations used for training, validation and testing are listed with their accession numbers in Supplementary Materials: Tables S1–S9. Assemblies and annotations used for the Mesangiospermae model that were not publicly released at the time of this study are available via Figshare (https://doi.org/10.6084/m9.figshare.32086728). Until their official release, their use is subject to the Fort Lauderdale Accord and users should contact the original data providers before use in independent analyses. The annotations of 2,948 vertebrate assemblies generated with Tiberius are available at https://bioinf.uni-greifswald.de/bioinf/tiberius/genes/tib-tbl-vert.html.

## 9 Author contributions statement

L.G. developed robust training instructions, performed quality control and generated benchmark comparisons, T.B. and S.T. trained plant parameters, A.Ka., A.Kr., A.S., I.V.G., and K.J.H. trained parameters for fungi, F.B., R.K. and L.G. improved Tiberius code, F.O., C.W.W., and K.J.H. trained Insecta parameters, M.S. made trainings and predictions for Vertebrata, K.J.H. trained Chlorophyta and Bacillariophyta parameters, K.J.H. developed the web server, M.S. and K.J.H. coordinated the project, L.G. wrote the first draft of the manuscript, all authors read and contributed to the final version of the manuscript.

## 10 Acknowledgments

We acknowledge EuroHPC Joint Undertaking for awarding us access to MareNostrum5 as BSC, Spain.

## 11 Supplementary Materials

Supplementary Figures, Methods and Tables. Contains the supplementary figures (Figures S1–S5) showing error patterns for the benchmarked tools, supplementary methods describing the Tiberius training procedure, accuracy metrics and benchmarking protocols for Tiberius, ANNEVO, Helixer and BRAKER3, followed by supplementary tables (Tables Sl-S32): Tables S1–S9 list genome assemblies, annotation sources and gene counts for all species used for training, validation and testing per clade; Tables S10–S16 provide per-species runtime benchmarks per clade; Table S17 documents the usage of Tiberius test species in the training sets of Helixer and ANNEVO; Tables S18–S25 provide per-species accuracy benchmarks per clade; Table S26 compares prediction accuracy between Tiberius v1.1.8 and v2.0.0; Table S27 compares runtimes between Tiberius v1.1.8 and v2.0.0; Table S28 lists the pre-trained models used for Helixer and ANNEVO in the benchmark; Tables S29–S30 describe hardware and hyperparameters used for model training and clade-specific model architectures; Table S31 reports paired Wilcoxon signed-rank tests comparing per-genome F1 scores of Tiberius with those of Helixer and ANNEVO; Table S32 gives the taxonomic distance and TimeTree divergence time between each test species and its closest training species.

## References

[1] Nuala A. O’Leary, Eric Cox, J. Bradley Holmes, W. Ray Anderson, Robert Falk, Vichet Hem, Mirian T. N. Tsuchiya, Gregory D. Schuler, Xuan Zhang, John Torcivia, Anne Keller. Laurie Breen, Jonathan Cothran, Hena Bajwa, Jovany Tinne, Peter A. Meric, Wratko Hlavina, and Valerie A. Schneider. Exploring and retrieving sequence and meta-data for species across the tree of life with NCBI Datasets. Scientific Data, 11(1):732, 2024. doi: 10.1038/s41597-024-03571-y.

[2] Mark Blaxter, Harris A Lewin, Federica DiPalma, Richard Challis, Manuela da Silva, Richard Durbin, Giulio Formenti, Nico Franz, Roderic Guigo, Peter W Harrison, Michael Hiller, Katharina J Hoff, Kerstin Howe, Erich D Jarvis, Mara K N Lawniczak, Kerstin Lindblad-Toh, Debra J H Mathews, Fergal J Martin, Camila J Mazzoni, Ann M McCartney, Nicola Mulder, Sadye Paez, Kim D Pruitt, Verena Ras, Oliver A Ryder, Lesley Shirley, Françoise Thibaud-Nissen, Tandy Warnow, Robert M Waterhouse, and EBP Community of Scientists. The Earth BioGenome Project Phase II: illuminating the eukaryotic tree of life. Frontiers in Science, 3:1514835, 2025. doi: 10.3389/fsci.2025.1514835.

[3] Caroline M. Weisman, Andrew W. Murray, and Sean R. Eddy. Mixing genome annotation methods in a comparative analysis inflates the apparent number of lineage-specific genes. Current Biology, 32(12):2632–2639, 2022. doi: 10.1016/j.cub.2022.04.085.

[4] L Gabriel, T Bruna, K J Hoff, M Ebel, A Lomsadze, M Borodovsky, and M Stanke. BRAKER3: Fully automated genome annotation using RNA-Seq and protein evidence with GeneMark-ETP, AUGUSTUS and TSEBRA. Genome Research, 34(5):769–777, 2024. doi: 10.1101/gr.278090.123.

[5] M Stanke and S Waack. Gene prediction with a hidden Markov model and a new intron submodel. Bioinformatics, 19(suppl_2):ii215–ii225, 2003. doi: 10.1093/bioinformatics/btgl080.

[6] Felix Holst, Anthony M Bolger, Felicitas Kindel, Janina Måls. Christopher Gunther, Sanjana Ayyar, Shweta Choudhari, Lifeng Wang, Andreas PM Weber, and Björn Usadel. Helixer: ab initio prediction of primary eukaryotic gene models combining deep learning and a hidden Markov model. Nature Methods, 22(11):2208–2217, 2025. doi: 10.1038/S41592-025-02939-1.

[7] Lars Gabriel, Felix Becker, Katharina J Hoff, and Mario Stanke. Tiberius: End-to-end deep learning with an HMM for gene prediction. Bioinformatics, 40(12):btae685, 2024. doi: 10.1093/bioinformatics/btae685.

[8] Pengyu Zhang, Tun Xu, Songbo Wang, Xiaofei Yang, Peisen Sun, Peng Jia, Jiadong Lin, Bo Wang, Yizhe Zhang, Deyu Meng, Stephen J Bush, Zemin Ning, and Kai Ye. Highly accurate ab initio gene annotation with ANNEVO. Nature Methods, 23:740–748, 2026. doi: 10.1038/s41592-026-03036-7.

[9] M Stanke, W Bruhn, F Becker, and KJ Hoff. VARUS: sampling complementary RNA reads from the sequence read archive. BMC Bioinformatics, 20:558, 2019. doi: 10.1186/S12859-019-3182-X.

[10] Rory J. Craig, Ahmed R. Hasan, Rob W. Ness, and Peter D. Keightley. Comparative genomics of Chlamydomonas. The Plant Cell, 33(4):1016–1041, 2021. doi: 10.1093/plcell/koab026.

[11] Gina V Filloramo, Bruce A Curtis, Emma Blanche, and John M Archibald. Reexamination of two diatom reference genomes using long-read sequencing. BMC Genomics, 22(1):379, 2021. doi: 10.1186/s12864-021-07666-3.

[12] Geo Pertea and Mihaela Pertea. GFF Utilities: GffRead and GffCompare. FlOOOResearch, 9:304, 2020. doi: 10.12688/f1000research.23297.2.

[13] Sudhir Kumar, Michael Suleski, Jack M Craig, Adrienne E Kasprowicz, Maxwell Sanderford, Michael Li, Glen Stecher, and S Blair Hedges. TimeTree 5: An expanded resource for species divergence times. Molecular Biology and Evolution, 39(8):msac174, 2022. doi: 10.1093/molbev/msacl74.

[14] The UniProt Consortium. UniProt: the universal protein knowledgebase in 2025. Nucleic Acids Research, 53(D1):D609–D617, 2025. doi: 10.1093/nar/gkael010.

[15] Benjamin Buchfink, Klaus Reuter, and Hajk-Georg Drost. Sensitive protein alignments at tree-of-life scale using DIAMOND. Nature Methods, 18(4):366–368, 2021. doi: 10.1038/s41592-021-01101-x.

[16] Tiberius vertebrate genome annotations (2,948 assemblies). https://bioinf.uni-greifswald.de/bioinf/tiberius/genes/tib-tbl-vert.html, 2026. Accessed 29 July 2026.

[17] Hiram Clawson, Brian T Lee, Brian J Raney, Galt P Barber, Jonathan Casper, Mark Diekhans, Clay Fischer, Jairo Navarro Gonzalez, Angie S Hinrichs, Christopher M Lee, Luis R Nassar, Gerardo Perez, Brittney Wick, Daniel Schmelter, Matthew L Speir, Joel Armstrong, Ann S Zweig, Robert M Kuhn, Bogdan M Kirilenko, Michael Hiller, David Haussler, W James Kent, and Maximilian Haeussler. GenArk: towards a million UCSC genome browsers. Genome Biology, 24(1):217, 2023. doi: 10.1186/s13059-023-03057-x.

[18] GenArk vertebrate genome assembly hub listing, https://hgdownload.gi.ucsc.edu/hubs/, 2023. Accessed 29 July 2026.

[19] Tiberius web server for eukaryotic genome annotation. https://bioinf.uni-greifswald.de/tiberius, 2025. Accessed 29 July 2026.

[20] Sean Whalen, Jacob Schreiber, William S. Noble, and Katherine S. Pollard. Navigating the pitfalls of applying machine learning in genomics. Nature Reviews Genetics, 23(3): 169–181, 2022. doi: 10.1038/s41576-021-00434-9.

[21] Stepan Saenko, Katharina J Hoff, and Mario Stanke. Annotation of 200 insect genomes with BRAKER for consistent comparisons across species. Scientific Data, 13(1):288, 2026. doi: 10.1038/s41597-026-06840-0.

[22] Natalia Nenasheva, Clara Pitzschel, Cynthia N Webster, Alexander J Hart, Jill L. Wegrzyn, Mia M Bengtsson, and Katharina J Hoff. Annotation of protein-coding genes in 49 diatom genomes from the Bacillariophyta clade. Scientific Data, 12(1):985, 2025. doi: 10.1038/s41597-025-05306-z.

[23] Moises Burset and Roderic Guigo. Evaluation of gene structure prediction programs. Genomics, 34(3):353–367, 1996. doi: 10.1006/geno.l996.0298.

[24] Genis Parra, Pankaj Agarwal, Josep F Abril, Thomas Wiehe, James W Fickett, and Roderic Guigó. Comparative gene prediction in human and mouse. Genome Research, 13(1):108–117, 2003. doi: 10.1101/gr.871403.

[25] Felipe A Simåo, Robert M Waterhouse, Panagiotis Ioannidis, Evgenia V Kriventseva, and Evgeny M Zdobnov. BUSCO: assessing genome assembly and annotation completeness with single-copy orthologs. Bioinformatics, 31 (19):3210–3212, 2015. doi: 10.1093/bioinformatics/btv351.

[26] Bogdan M Kirilenko, Chetan Munegowda, Ekaterina Osipova, David Jebb, Virag Sharma, Moritz Blumer, Ariadna E Morales, Alexis-Walid Ahmed, Dimitrios-Georgios Kontopoulos, Leon Hilgers, Kerstin Lindblad-Toh, Elinor K Karlsson, Zoonomia Consortium, and Michael Hiller. Integrating gene annotation with orthology inference at scale. Science, 380(6643):eabn3107, 2023. doi: 10.1126/science.abn3107.

[27] Corentin Meyer, Nicolas Scalzitti, Anne Jeannin-Girardon, Pierre Collet, Olivier Poch, and Julie D Thompson. Understanding the causes of errors in eukaryotic protein-coding gene prediction: a case study of primate proteomes. BMC Bioinformatics, 21(1):513, 2020. doi: 10.1186/s12859-020-03855-l.

