## Supplementary Materials for "Accurate ab *initio* gene prediction in eukaryotes with Tiberius in multiple clades"

Lars Gabriel[1], Tomáš Brůna[2], Asees Kaur[3], Anish Krishnan[4], Felix Ortmann[1], Asaf Salamov[2], Samuel Talbot[5], Felix Becker[1], Richard Krieg[1], Christopher W. Wheat[6], Igor V. Grigoriev[2,4], Mario Stanke[1,*], and Katharina J. Hoff[1,*]

[1]Institute of Mathematics and Computer Science, University of Greifswald, Walther-Rathenau-Str. 47, 17489 Greifswald, Germany
[2]DOE Joint Genome Institute, Lawrence Berkeley National Laboratory, 1 Cyclotron Road, Berkeley, 94720, California, USA
[3]University of California Merced, Merced, CA 95343, USA
[4]Department of Plant and Microbial Biology, University of California Berkeley, Berkeley, CA 94720, USA
[5]Center for Quantitative Life Sciences, Oregon State University, OR 97331, USA
[6]Department of Zoology, Stockholm University, Sweden

#### Supplementary Figures

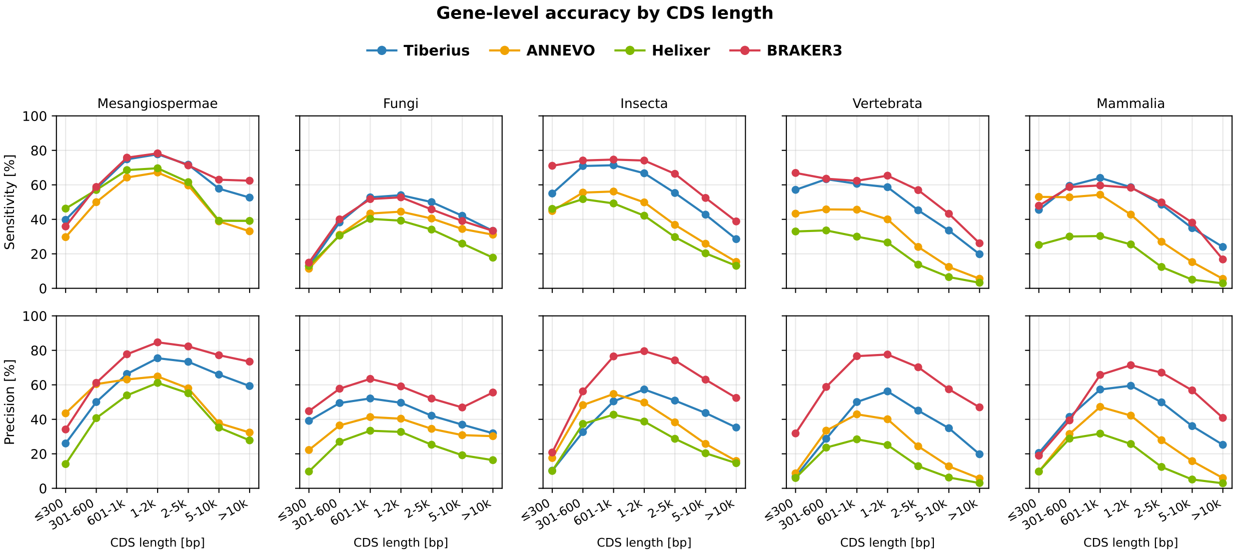


Figure S1: Gene-level accuracy of Tiberius, ANNEVO, Helixer and BRAKER3 stratified by coding sequence (CDS) length. For each test genome, reference genes and predicted genes were separately assigned to CDS-length bins (x-axis) and `gffcompare` was run once per bin, comparing the predicted genes of that bin with the reference genes of the same bin. A prediction that matches a reference gene exactly has the same CDS and therefore falls into the same bin. Upper row: sensitivity, the percentage of reference genes in a bin that were predicted correctly. Lower row: precision, the percentage of predicted genes in a bin that exactly match a reference gene. Low precision in a bin thus indicates many predictions of that length without a reference counterpart, i.e. overpredicted or incorrectly structured genes. Columns show clades; Vertebrata denotes non-mammalian vertebrates.

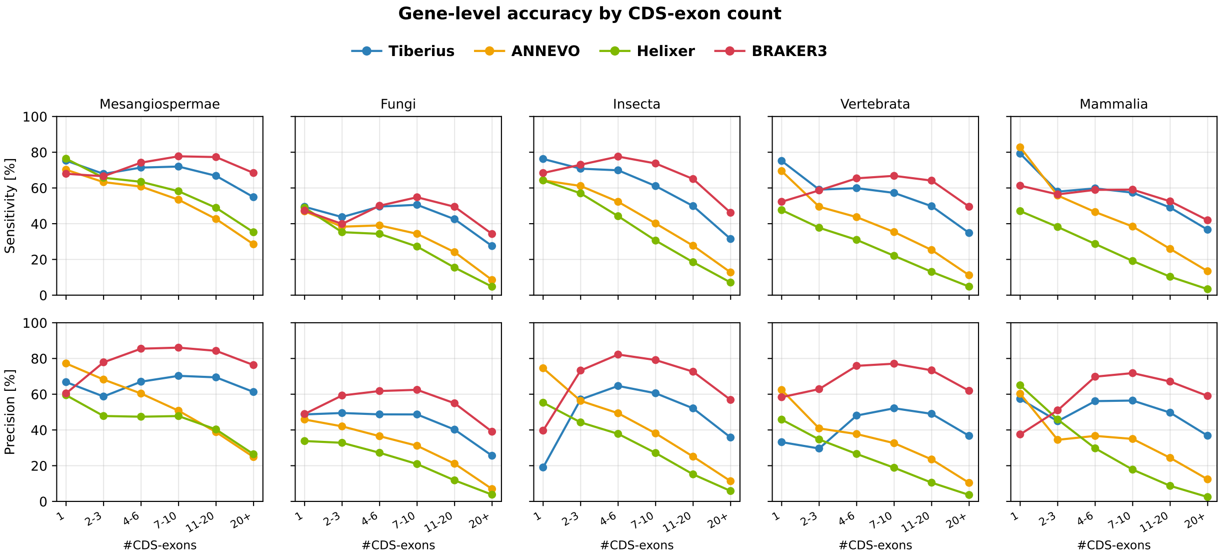


Figure S2: Gene-level accuracy of Tiberius, ANNEVO, Helixer and BRAKER3 stratified by the number of CDS exons per gene. For each test genome, reference genes and predicted genes were separately assigned to CDS-exon-count bins (x-axis) and `gffcompare` was run once per bin, comparing the predicted genes of that bin with the reference genes of the same bin. A prediction that matches a reference gene exactly has the same number of CDS exons and therefore falls into the same bin. Upper row: sensitivity, the percentage of reference genes in a bin that were predicted correctly. Lower row: precision, the percentage of predicted genes in a bin that exactly match a reference gene. Low precision in a bin thus indicates many predictions with that number of CDS exons without a reference counterpart, i.e. overpredicted or incorrectly structured genes. Columns show clades; Vertebrata denotes non-mammalian vertebrates.

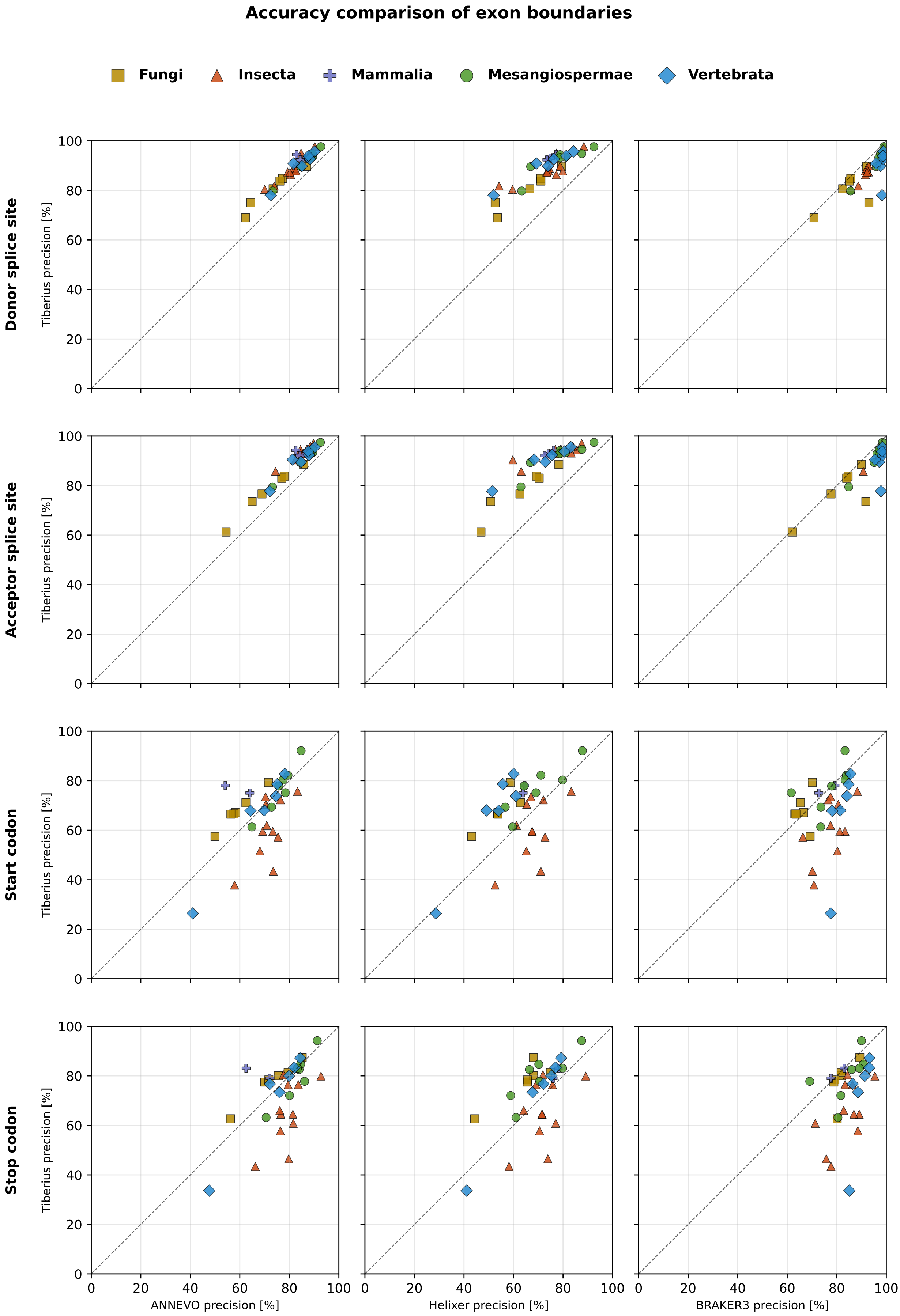


Figure S3: Per-species scatter comparing Tiberius against ANNEVO, Helixer and BRAKER3 on the precision of four coding-exon boundary types: donor and acceptor splice sites and start and stop codons. Each dot is one benchmark species, coloured by clade. The dashed line marks equal precision, points above it indicate Tiberius matches the reference more often than the compared tool for the boundary type.

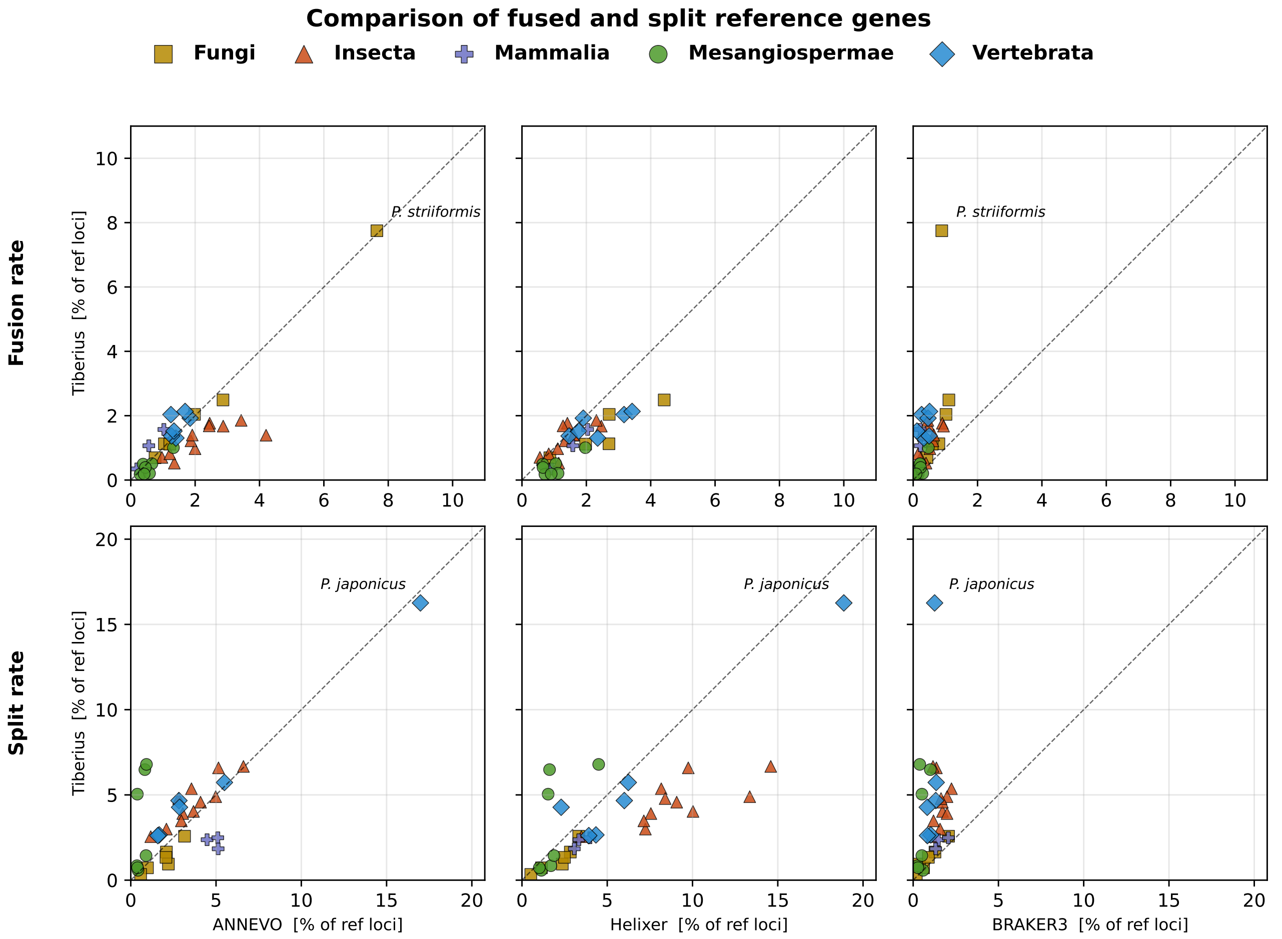


Figure S4: Per-species scatter comparing Tiberius against ANNEVO, Helixer and BRAKER3 on the fraction of reference gene loci that were affected by fusion and split errors. Each dot is one benchmark species, coloured by clade. The dashed line marks equal error, points below it indicate Tiberius makes fewer errors than the compared tool. Gene loci were defined separately for the reference and for each tool's prediction: transcripts on the same strand whose genomic spans (from the start of the first exon to the end of the last exon) overlap were grouped into one locus, and each locus is represented by its longest isoform. A reference locus and a predicted locus are considered matched if their coding sequences overlap by at least 30 nt on the same strand and in the same reading frame. Fusion rate is the fraction of reference loci whose matched predicted locus is also matched to at least one other reference locus. Split rate is the fraction of reference loci matched to more than one predicted locus.

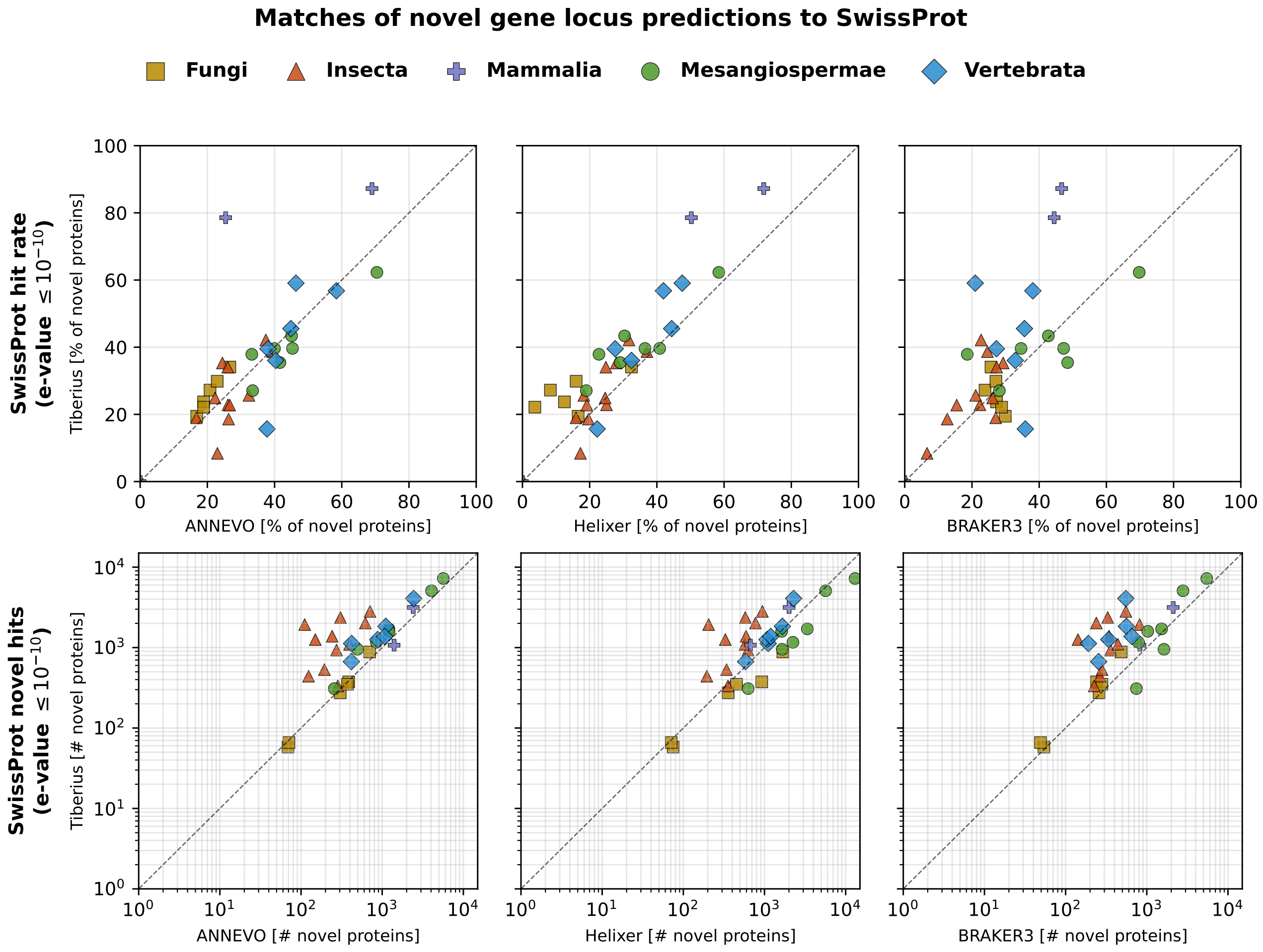


Figure S5: For each predicted gene locus with no in-frame CDS overlap ($\geq 30$ nt, same strand) with any reference locus, we searched the resulting protein against UniProt SwissProt release 2026_03 [1] with DIAMOND blastp (version 2.0.14) [2]. Per-species scatter comparing Tiberius against ANNEVO, Helixer, and BRAKER3. Row 1 shows the SwissProt hit rate, the percentage of a tool's novel loci per species whose protein has a hit at e-value $\leq 10^{-10}$. Row 2 shows the absolute number of novel loci with SwissProt homology (log scale). Points above the diagonal indicate that Tiberius has a higher hit rate (row 1) or more novel loci with SwissProt homology (row 2) than the compared tool.

### Supplementary Methods

#### Tiberius Training

The Tiberius models were trained as described here: `https://github.com/Gaius-Augustus/Tiberius/blob/main/docs/training_large_data.md`, commit `46d7bc4`.

#### Accuracy metrics

Accuracy metrics were computed using `gffcompare` v0.12.10 on the basis of CDS features. Gffcompare was used so that instances from a prediction `prediction.gff` were only counted as true positives if they matched the reference `reference.gff` exactly. A tolerance of 3 bp was given to allow for differences in stop codon inclusion in the CDS features of the files.

```
gffcompare --strict-match -e 3 -T -r reference.gff prediction.gff
```

#### Benchmarking Tiberius

Tiberius v2.0.0 was used with one of the pretrained models for each test species, see Table S28. For a target genome `genome.fa` and a Tiberius model `${model}`, Tiberius was run with:

```
tiberius.py --genome genome.fa --model_cfg ${model}.yaml --out tiberius.gtf
```

#### Benchmarking ANNEVO

ANNEVO v2.2.1 was used with an appropriate pretrained model for each test species, see Table S28. For a target genome `genome.fa` and an ANNEVO model `${model}.pt`, ANNEVO was run with:

```
annotation.py --genome genome.fa --model_path ${model}.pt --threads 72 --output annevo.gff
```

#### Benchmarking Helixer

Helixer v0.3.4 was used with an appropriate pretrained model for each test species, see Table S28. For a target genome `genome.fa` and a Helixer model `${model}`, Helixer was run with:

```
Helixer.py --fasta-path genome.fa --lineage ${model} --gff-output-path helixer.gff
```

#### Benchmarking BRAKER3

BRAKER3 v3.0.8 was run using a softmasked genome, protein sequences and aligned RNA-Seq short reads as inputs. The proteomes of all training species from the Tiberius training of the clade of the target genome were used as the protein input `proteins.fa`.

The unmasked input genome `genome_dir/genome.fa` was softmasked for repeats using RED v2.0 with:

```
Red -gnm genome_dir/ -msk red/ -cor ${params.threads}
```

The short-read RNA-Seq input was generated with VARUS, which computes a `BAM` file with aligned reads that have sufficient coverage of the genome for genome annotation (resulting in `VARUS.bam`).

```
perl runVARUS.pl \
    --readFromTable=0 \
    --createindex=1 \
    --latinGenus=$genus_name \
    --latinSpecies=$species_name \
    --speciesGenome=genome.red.fa \
    --aligner=HISAT \
```

```
    --runThreadN 72 \
    --batchSize 50000 \
    --blockSize 5000 \
    --maxBatches 1000 \
    --mergeThreshold 10
```

BRAKER3 was then run with:

```
braker.pl \
    --genome=genome.red.fa \
    --prot_seq=proteins.fa \
    --threads=72 \
    --workingdir=braker3 \
    --bam VARUS.bam
```

### Supplementary Tables

| Species | Accession ID | Genome (Mb) | Annotation | #Genes | Species | Accession ID | Genome (Mb) | Annotation | #Genes |
|---|---|---|---|---|---|---|---|---|---|
| **Training** | | | | | | | | | |
| *Cyclotella cryptica* | GCA_013187285.1 | 171 | BRAKER | 16,338 | *Cyclotella choctawhatcheeana* | GCA_036939855.1 | 55 | BRAKER | 15,687 |
| *Conticribra guillardii* | GCA_036939335.1 | 75 | BRAKER | 15,775 | *Conticribra weissflogii* | GCA_036940025.1 | 130 | BRAKER | 14,524 |
| *Skeletonema menzelii* | GCA_036940005.1 | 33 | BRAKER | 13,949 | *Skeletonema potamos* | GCA_036940105.1 | 35 | BRAKER | 13,943 |
| *Discostella pseudostelligera* | GCA_036940085.1 | 30 | BRAKER | 10,458 | *Discostella stelligera* | GCA_036939735.1 | 59 | BRAKER | 13,154 |
| *Stephanodiscus triporus* | GCA_036939755.1 | 58 | BRAKER | 13,345 | *Cyclostephanos invisitatus* | GCA_036939675.1 | 58 | BRAKER | 12,825 |
| *Thalassiosira mediterranea* | GCA_036939795.1 | 77 | BRAKER | 16,656 | *Cylindrotheca closterium* | GCA_933822405.4 | 55.1 | EMBL | 23,336 |
| *Pseudo-nitzschia multistriata* | GCA_900660405.1 | 57 | EMBL | 11,733 | *Mayamaea pseudoterrestris* | GCA_027923505.1 | 31 | DDBJ | 10,842 |
| *Seminavis robusta* | GCA_903772945.1 | 126 | EMBL | 34,364 | *Fragilaria crotonensis* | GCA_022925895.1 | 62 | Genbank | 25,842 |
| *Asterionella formosa* | GCA_002256025.1 | 68 | BRAKER | 14,820 | *Chaetoceros muelleri* | GCA_019693545.1 | 38 | BRAKER | 12,172 |
| *Chaetoceros tenuissimus* | GCA_021927905.1 | 41 | DDBJ | 18,636 | | | | | |
| **Validation** | | | | | | | | | |
| *Nitzschia palea* | GCA_019593585.1 | 41 | BRAKER | 15,386 | *Skeletonema tropicum* | GCA_037178625.1 | 79 | BRAKER | 23,802 |
| **Testing** | | | | | | | | | |
| *Thalassiosira pseudonana* | Filloramo *et al.* [3] | 34 | BRAKER2/PASA | 14,830 | *Phaeodactylum tricornutum* | GCF_000150955.2 | 27.6 | ENSEMBL | 11,909 |

Table S1: Accession ID, genome size, annotation source, and gene count for all species used for training, validation, and testing the Tiberius model for **Bacillariophyta**. The BRAKER annotations were provided by Nenasheva *et al.* [4], the genome and annotation of *Thalassiosira pseudonana* was downloaded from `https://doi.org/10.5683/SP2/ZDZQFE` [3]; this annotation was generated by Filloramo *et al.* with a pipeline based on BRAKER2 and subsequently corrected with PASA.

| Species | Accession ID | Genome (Mb) | Annotation | #Genes | Species | Accession ID | Genome (Mb) | Annotation | #Genes |
|---|---|---|---|---|---|---|---|---|---|
| **Training** | | | | | | | | | |
| *Ostreococcus lucimarinus* CCE9901 | GCF_000092065.1 | 13 | RefSeq | 6,381 | *Chlamydomonas reinhardtii* | GCF_000002595.2 | 111.1 | RefSeq | 17,705 |
| *Volvox carteri f. nagariensis* | GCF_000143455.1 | 138 | RefSeq | 12,362 | *Picochlorum sp.* BPE2 | GCA_025209345.1 | 14.9 | Genbank | 7,227 |
| *Astrephomene gubernaculifera* | GCA_021605115.1 | 104 | DDBJ | 7,209 | *Nannochloris sp. desiccata* | GCA_019044685.2 | 21.6 | Genbank | 9,191 |
| *Dunaliella primolecta* | GCA_914767535.2 | 211 | ensembl | 10,240 | *Chloropicon primus* | GCA_023205875.1 | 17.6 | Genbank | 8,513 |
| *Tetradesmus obliquus* | GCA_030272155.1 | 105 | Genbank | 14,724 | | | | | |
| **Validation** | | | | | | | | | |
| *Micromonas commoda* | GCF_000090985.2 | 21 | RefSeq | 9,201 | *Gonium pectorale* | GCA_001584585.1 | 149 | Genbank | 16,290 |
| **Testing** | | | | | | | | | |
| *Edaphochlamys debaryana* | GCA_016858145.1 | 142 | Genbank | 19,013 | *Bathycoccus prasinos* | GCF_002220235.1 | 15 | RefSeq | 7,813 |

Table S2: Accession ID, genome size, annotation source, and gene count for all species used for training, validation, and testing the Tiberius model for **Chlorophyta**. The GenBank annotation of the test species *Edaphochlamys debaryana* was generated with BRAKER v2.1.2 by Craig *et al.* [5].

| Species | Accession ID | Genome (Mb) | #Genes | Annotation source | Reference | DOI |
|---|---|---|---|---|---|---|
| **Training** | | | | | | |
| *Acorus americanus* | 586 | 374 | 26,508 | Phytozome | Leebens-Mack et al. (2017) | doi:10.46936/10.25585/60001405 |
| *Agave tequilana* | 924 | 3,394 | 42,295 | Phytozome | Xiaohan et al. (2017) | doi:10.46936/10.25585/60001139 |
| *Aquilegia coerulea* | 322 | 292 | 30,023 | Phytozome | Filiault et al. (2018) | doi:10.7554/eLife.36426 |
| *Arundo donax* | 1047[1] | 474 | 25,466 | Phytozome | Blake et al. (2019) | doi:10.46936/10.25585/60001226 |
| *Beta vulgaris* | 782 | 535 | 21,587 | Phytozome | McGrath et al. (2023) | doi:10.1093/dnares/dsac033 |
| *Brassica rapa* | 714 | 432 | 39,062 | Phytozome | Greenham et al. (2018) | doi:10.46936/10.25585/60001202 |
| *Brocchinia micrantha* | 933[1] | 338 | 20,842 | Phytozome | Leebens-Mack et al. (2017) | doi:10.46936/10.25585/60001405 |
| *Capparis spinosa* | 851[1] | 575 | 25,776 | Phytozome | Harkess et al. (2020) | doi:10.46936/10.25585/60000482 |
| *Chasmanthium laxum* | 690 | 863 | 31,776 | Phytozome | Schnable et al. (2013) | doi:10.46936/10.25585/60001029 |
| *Citrus limon* | 831[1] | 321 | 25,726 | Phytozome | - | - |
| *Cleome violacea* | 585 | 279 | 21,850 | Phytozome | Wing et al. (2011) | doi:10.46936/10.25585/60000980 |
| *Coffea arabica* | 871 | 1,025 | 51,583 | Phytozome | Medrano et al. (2024) | doi:10.1093/g3journal/jkae262 |
| *Corylus avellana* | 858 | 350 | 32,431 | Phytozome | Talbot et al. (2024) | doi:10.1093/g3journal/jkae021 |
| *Daucus carota* | 861 | 441 | 36,307 | Phytozome | Coe et al. (2023) | doi:10.1038/s41477-023-01526-6 |
| *Dendrobium catenatum* | GCA_002786265.1 | 1,061 | 22,642 | NCBI | Zhang et al. (2017) | doi:10.1038/nature23897 |
| *Dioscorea alata* | 550 | 480 | 25,189 | Phytozome | Bredeson et al. (2022) | doi:10.1038/s41467-022-29114-w |
| *Eucalyptus grandis* | 891 | 571 | 35,929 | Phytozome | Lötter et al. (2025) | doi:10.1093/g3journal/jkaf112 |
| *Fragaria vesca* | 677 | 219 | 34,006 | Phytozome | Li et al. (2019) | doi:10.1038/s41438-019-0142-6 |
| *Glycine max* | 880 | 1,011 | 48,387 | Phytozome | Espina et al. (2024) | doi:10.1111/tpj.17026 |
| *Gossypium hirsutum* | 695 | 2,296 | 75,366 | Phytozome | - | - |
| *Hydrocotyle leucocephala* | 768 | 756 | 32,364 | Phytozome | Bertrand et al. (2019) | doi:10.46936/10.25585/60001311 |
| *Joinvillea ascendens* | 587 | 1,213 | 29,122 | Phytozome | Leebens-Mack et al. (2017) | doi:10.46936/10.25585/60001405 |
| *Lactuca sativa* | LsPKU06 | 2,593 | 45,981 | Figshare | Wang et al. (2024) | doi:10.1016/j.xplc.2024.101011 |
| *Lindenbergia philippensis* | 689 | 426 | 26,721 | Phytozome | Leebens-Mack et al. (2017) | doi:10.46936/10.25585/60001405 |
| *Malus domestica* | WA 38 v1.0 | 652 | 54,235 | Genome Database for Rosaceae | Zhang et al. (2024) | doi:10.1093/g3journal/jkae222 |
| *Manihot esculenta* | 671 | 637 | 32,805 | Phytozome | Bredeson et al. (in prep.) | - |
| *Mentha longifolia* | -[1] | 418 | 42,275 | Oregon State University | Talbot et al. (in prep.) | - |
| *Miscanthus sinensis* | 497 | 1,847 | 67,789 | Phytozome | Mitros et al. (2020) | doi:10.1038/s41467-020-18923-6 |
| *Moringa oleifera* | 866[1] | 271 | 20,761 | Phytozome | Harkess et al. (2020) | doi:10.46936/10.25585/60000482 |
| *Musa acuminata* | GCF_036884655.1 | 477 | 29,948 | NCBI | Huang et al. (2023) | doi:10.1093/hr/uhad153 |
| *Myrothamnus flabellifolia* | 856 | 1,283 | 22,809 | Phytozome | Marks et al. (2025) | doi:10.1111/nph.70700 |
| *Nicotiana tabacum* | 945 | 4,322 | 66,812 | Phytozome | Wang et al. (2024) | doi:10.1016/j.molp.2024.01.008 |
| *Oryza sativa* | 499 | 378 | 35,594 | Phytozome | Jain et al. (2019) | doi:10.1186/s12864-019-6262-4 |
| *Osyris compressa* | 823[1] | 278 | 29,654 | Phytozome | Leebens-Mack et al. (2017) | doi:10.46936/10.25585/60001405 |
| *Panicum virgatum* | 516 | 527 | 40,957 | Phytozome | Lovell et al. (2021) | doi:10.1038/s41586-020-03127-1 |
| *Pharus latifolius* | 680 | 1,118 | 32,144 | Phytozome | Leebens-Mack et al. (2017) | doi:10.46936/10.25585/60001405 |
| *Phoenix dactylifera* | GCA_009389715.1 | 773 | 29,236 | NCBI | Hazzouri et al. (2019) | doi:10.1038/s41467-019-12604-9 |
| *Setaria viridis* | 726 | 397 | 29,807 | Phytozome | Mamidi et al. (2020) | doi:10.1038/s41587-020-0681-2 |
| *Sorghum bicolor* | 730 | 720 | 32,160 | Phytozome | Morris et al. (2025) | doi:10.1101/2025.08.01.667986 |
| *Thinopyrum intermedium* | 770 | 4,303 | 41,523 | Phytozome | Poland et al. (2015) | doi:10.46936/10.25585/60000844 |
| *Ulmus americana* | 881 | 1,461 | 25,621 | Phytozome | - | - |
| *Vitis vinifera* | -[1] | 488 | 40,588 | Oregon State University | Talbot et al. (in prep.) | - |
| *Yucca filamentosa* | 837 | 2,422 | 45,250 | Phytozome | Heyduk et al. (2016) | doi:10.46936/10.25585/60001082 |
| *Zea mays* | 833 | 2,178 | 39,756 | Phytozome | Hufford et al. (2021) | doi:10.1126/science.abg5289 |
| *Zingiber officinale* | GCF_018446385.1 | 1,625 | 33,352 | NCBI | - | - |
| **Validation** | | | | | | |
| *Populus trichocarpa* | 533 | 392 | 34,699 | Phytozome | Tuskan et al. (2006) | doi:10.1126/science.1128691 |
| *Typha latifolia* | 853 | 215 | 22,107 | Phytozome | Leebens-Mack et al. (2017) | doi:10.46936/10.25585/60001405 |
| **Testing** | | | | | | |
| *Arabidopsis thaliana* | 447 | 119 | 27,655 | Phytozome | Cheng et al. (2017) | doi:10.1111/tpj.13415 |
| *Brachypodium distachyon* | GCF_000005505.3 | 271 | 25,534 | NCBI | Vogel et al. (2010) | doi:10.1038/nature08747 |
| *Eschscholzia californica* | 828 | 384 | 29,637 | Phytozome | Rössner et al. (2026) | doi:10.1093/plcell/koag039 |
| *Freycinetia multiflora* | 868 | 511 | 28,350 | Phytozome | Leebens-Mack et al. (2017) | doi:10.46936/10.25585/60001405 |
| *Medicago truncatula* | GCF_003473485.1 | 429 | 31,925 | NCBI | Pecrix et al. (2018) | doi:10.1038/s41477-018-0286-7 |
| *Mimulus guttatus* | 797 | 339 | 25,113 | Phytozome | Lovell et al. (2025) | doi:10.1111/1755-0998.70012 |
| *Urochloa brizantha* | 897[1] | 2,747 | 134,704 | Phytozome | Ozias-Akins et al. (2020) | doi:10.46936/10.25585/60001353 |

Table S3: Genome assemblies and annotation sources used for training, validation and testing the **Mesangiospermae** model.

[1]Not yet publicly released at the time of this study; used with permission from the data providers. All genomes and annotations used for training are available via Figshare (doi:10.6084/m9.figshare.32086728) for reproducibility purposes. Until their official public release on Phytozome or elsewhere, use of these data is subject to the Fort Lauderdale Accord and users should contact the original data providers before use in independent analyses.

| Species | Accession ID | Genome (Mb) | #Genes | Species | Accession ID | Genome (Mb) | #Genes |
|---|---|---|---|---|---|---|---|
| **Training (1/2)** | | | | | | | |
| *Aaosphaeria arxii* | GCF_010015735.1 | 39 | 14,293 | *Akanthomyces muscarius* | GCF_028009165.1 | 36 | 12,347 |
| *Alternaria atra* | GCF_907166805.1 | 40 | 12,172 | *Alternaria burnsii* | GCF_013036055.1 | 33 | 11,314 |
| *Alternaria rosae* | GCF_020736505.1 | 34 | 12,727 | *Amorphotheca resinae* | GCF_003019875.1 | 29 | 9,642 |
| *Annulohypoxylon maeteangense* | GCF_022496945.1 | 38 | 11,144 | *Annulohypoxylon truncatum* | GCF_022578515.1 | 38 | 11,304 |
| *Apiospora aurea* | GCF_038362705.1 | 50 | 15,187 | *Apiospora marii* | GCF_038362585.1 | 51 | 16,453 |
| *Apiotrichum porosum* | GCF_003942205.1 | 25 | 9,719 | *Aplosporella prunicola* | GCF_010093885.1 | 33 | 12,738 |
| *Ascochyta rabiei* | GCF_004011695.2 | 41 | 11,642 | *Aspergillus aculeatus* | GCF_001890905.1 | 35 | 11,152 |
| *Aspergillus alliaceus* | GCF_009176365.1 | 40 | 13,336 | *Aspergillus brunneoviolaceus* | GCF_003184695.1 | 37 | 12,377 |
| *Aspergillus caelatus* | GCF_009193585.1 | 40 | 14,157 | *Aspergillus campestris* | GCF_002847485.1 | 28 | 9,823 |
| *Aspergillus candidus* | GCF_002847045.1 | 27 | 9,803 | *Aspergillus chevalieri* | GCF_016861735.1 | 30 | 10,516 |
| *Aspergillus clavatus* | GCF_000002715.2 | 28 | 9,375 | *Aspergillus fijiensis* | GCF_003184825.1 | 37 | 12,324 |
| *Aspergillus fischeri* | GCF_000149645.3 | 31 | 10,664 | *Aspergillus glaucus* | GCF_001890805.1 | 28 | 11,428 |
| *Aspergillus heteromorphus* | GCF_003184545.1 | 36 | 11,088 | *Aspergillus homomorphus* | GCF_003184865.1 | 34 | 11,651 |
| *Aspergillus ibericus* | GCF_003184845.1 | 33 | 11,983 | *Aspergillus japonicus* | GCF_003184785.1 | 36 | 12,338 |
| *Aspergillus lentulus* | GCF_010724455.1 | 30 | 10,506 | *Aspergillus neoniger* | GCF_003184625.1 | 35 | 12,218 |
| *Aspergillus piperis* | GCF_003184755.1 | 35 | 12,357 | *Aspergillus puulaauensis* | GCF_016861865.1 | 34 | 13,748 |
| *Aspergillus saccharolyticus* | GCF_003184585.1 | 31 | 10,375 | *Aspergillus sclerotioniger* | GCF_003184525.1 | 37 | 12,603 |
| *Aspergillus steynii* | GCF_002849105.1 | 38 | 13,146 | *Aspergillus sydowii* | GCF_001890705.1 | 34 | 13,716 |
| *Aspergillus tubingensis* | GCF_013340325.1 | 35 | 11,809 | *Aspergillus udagawae* | GCF_001078395.1 | 32 | 11,009 |
| *Aspergillus uvarum* | GCF_003184745.1 | 36 | 12,342 | *Aspergillus vadensis* | GCF_003184925.1 | 36 | 12,402 |
| *Aspergillus versicolor* | GCF_001890125.1 | 33 | 13,361 | *Aspergillus viridinutans* | GCF_018404265.1 | 35 | 10,237 |
| *Aspergillus wentii* | GCF_001890725.1 | 31 | 12,657 | *Aureobasidium melanogenum* | GCF_000721775.1 | 26 | 10,582 |
| *Aureobasidium namibiae* | GCF_000721765.1 | 25 | 10,259 | *Aureobasidium pullulans* | GCF_000721785.1 | 30 | 11,844 |
| *Babjeviella inositovora* | GCF_001661335.1 | 15 | 6,795 | *Batrachochytrium dendrobatidis* | GCF_000203795.2 | 24 | 8,386 |
| *Beauveria bassiana* | GCF_000280675.1 | 34 | 10,364 | *Bipolaris oryzae* | GCF_000523455.1 | 31 | 12,000 |
| *Bipolaris victoriae* | GCF_000527765.1 | 33 | 12,882 | *Bipolaris zeicola* | GCF_000523435.1 | 31 | 12,848 |
| *Boeremia exigua* | GCF_020726555.1 | 35 | 12,996 | *Botrytis byssoidea* | GCF_014898295.1 | 43 | 12,204 |
| *Botrytis cinerea* | GCF_000143535.2 | 43 | 11,698 | *Botrytis deweyae* | GCF_014898535.1 | 44 | 12,465 |
| *Botrytis fragariae* | GCF_013461495.1 | 42 | 12,345 | *Botrytis porri* | GCF_014898465.1 | 47 | 12,076 |
| *Botrytis sinoallii* | GCF_014898435.1 | 61 | 12,200 | *Canariomyces notabilis* | GCF_033297395.1 | 31 | 11,102 |
| *Candida albicans* | GCF_000182965.3 | 14 | 6,263 | *Candida jiufengensis* | GCF_024610255.1 | 14 | 5,705 |
| *Candida orthopsilosis* | GCF_000315875.1 | 13 | 5,782 | *Candida parapsilosis* | GCF_000182765.1 | 13 | 5,950 |
| *Capronia coronata* | GCF_000585585.1 | 26 | 9,231 | *Capronia epimyces* | GCF_000585565.1 | 29 | 10,468 |
| *Cercospora kikuchii* | GCF_019650295.1 | 34 | 13,001 | *Chaetomium fimeti* | GCF_033439085.1 | 35 | 11,628 |
| *Chaetomium strumarium* | GCF_033439665.1 | 32 | 10,401 | *Chaetomium tenue* | GCF_020726465.1 | 34 | 11,734 |
| *Cladophialophora bantiana* | GCF_000835475.1 | 37 | 12,817 | *Cladophialophora carrionii* | GCF_000365165.1 | 29 | 10,428 |
| *Cladophialophora psammophila* | GCF_000585535.1 | 39 | 13,421 | *Cladophialophora yegresii* | GCF_000585515.1 | 28 | 10,118 |
| *Clavispora lusitaniae* | GCF_014636115.1 | 12 | 5,702 | *Coccidioides immitis* | GCF_000149335.2 | 29 | 9,905 |
| *Cokeromyces recurvatus* | GCF_025118155.1 | 28 | 11,066 | *Colletotrichum acutatum* | GCF_030867785.1 | 49 | 15,371 |
| *Colletotrichum godetiae* | GCF_030913485.1 | 52 | 16,406 | *Colletotrichum graminicola* | GCF_000149035.1 | 52 | 12,399 |
| *Colletotrichum karsti* | GCF_011947395.1 | 52 | 13,328 | *Colletotrichum navitas* | GCF_030913465.1 | 52 | 14,963 |
| *Colletotrichum orchidophilum* | GCF_001831195.1 | 49 | 14,452 | *Colletotrichum paranaense* | GCF_030867605.1 | 49 | 16,727 |
| *Colletotrichum phormii* | GCF_030913505.1 | 52 | 15,598 | *Colletotrichum tamarilloi* | GCF_030869305.1 | 52 | 17,330 |
| *Coniosporium apollinis* | GCF_000281105.1 | 29 | 9,367 | *Cryptococcus amylolentus* | GCF_001720205.1 | 20 | 8,503 |
| *Cryptococcus bacillisporus* | GCF_000836335.1 | 18 | 6,809 | *Cryptococcus decagattii* | GCF_036417295.1 | 18 | 6,671 |
| *Cryptococcus deuterogattii* | GCF_002954075.1 | 18 | 6,673 | *Cryptococcus gattii* | GCF_000185945.1 | 18 | 6,580 |
| *Cryptococcus neoformans* | GCF_000091045.1 | 19 | 7,004 | *Cryptococcus tetragattii* | GCF_000835755.1 | 18 | 6,807 |
| *Cryptococcus wingfieldii* | GCF_001720155.1 | 20 | 8,314 | *Cucurbitaria berberidis* | GCF_010015615.1 | 33 | 12,468 |
| *Cyberlindnera jadinii* | GCF_001661405.1 | 13 | 6,184 | *Cyphellophora europaea* | GCF_000365145.1 | 29 | 11,153 |
| *Cystobasidium minutum* | GCF_039999085.1 | 21 | 7,862 | *Daldinia caldariorum* | GCF_022478825.1 | 36 | 10,553 |
| *Daldinia decipiens* | GCF_022478715.1 | 35 | 10,881 | *Daldinia vernicosa* | GCF_022497035.1 | 36 | 10,827 |
| *Debaryomyces fabryi* | GCF_001447935.2 | 12 | 6,025 | *Debaryomyces hansenii* | GCF_000006445.2 | 12 | 6,520 |
| *Desarmillaria tabescens* | GCF_030435595.1 | 75 | 19,381 | *Diaporthe amygdali* | GCF_026229845.1 | 51 | 15,635 |
| *Dichomitus squalens* | GCF_000275845.1 | 43 | 12,497 | *Dichotomopilus funicola* | GCF_033296635.1 | 33 | 9,442 |
| *Didymella exigua* | GCF_010094145.1 | 34 | 12,508 | *Dioszegia hungarica* | GCF_025882075.1 | 21 | 8,321 |
| *Diplodia corticola* | GCF_001883845.1 | 35 | 10,831 | *Dipodascopsis tothii* | GCF_038497935.1 | 13 | 5,897 |
| *Dothidotthia symphoricarpi* | GCF_010015815.1 | 34 | 11,821 | *Drechmeria coniospora* | GCF_001625195.1 | 33 | 8,445 |
| *Drepanopeziza brunnea* | GCF_022702325.1 | 52 | 9,615 | *Durotheca rogersii* | GCF_024521655.1 | 48 | 10,623 |
| *Emericellopsis atlantica* | GCF_019669845.1 | 27 | 10,066 | *Eremothecium gossypii* | GCF_000091025.4 | 9 | 5,317 |
| *Eremothecium sinecaudum* | GCF_001548555.1 | 9 | 4,970 | *Exophiala aquamarina* | GCF_000709125.1 | 42 | 13,118 |
| *Exophiala dermatitidis* | GCF_000230625.1 | 26 | 9,355 | *Exophiala oligosperma* | GCF_000835515.1 | 38 | 11,938 |
| *Exophiala spinifera* | GCF_000836115.1 | 33 | 12,110 | *Exophiala viscosa* | GCF_022695815.1 | 28 | 11,369 |
| *Exophiala xenobiotica* | GCF_000835505.1 | 31 | 12,077 | *Fimicolochytrium jonesii* | GCF_025526975.1 | 31 | 10,236 |
| *Fomitiporia mediterranea* | GCF_000271605.1 | 63 | 11,411 | *Fonsecaea erecta* | GCF_001651985.1 | 35 | 12,120 |
| *Fonsecaea multimorphosa* | GCF_000836435.1 | 33 | 12,405 | *Fulvia fulva* | GCF_020509005.1 | 67 | 14,993 |
| *Fusarium fujikuroi* | GCF_900079805.1 | 44 | 14,943 | *Fusarium mangiferae* | GCF_900044065.1 | 46 | 16,140 |
| *Fusarium odoratissimum* | GCF_000260195.1 | 47 | 16,975 | *Fusarium proliferatum* | GCF_900067095.1 | 45 | 16,488 |
| *Fusarium pseudograminearum* | GCF_000303195.2 | 37 | 12,445 | *Fusarium redolens* | GCF_020744475.1 | 53 | 17,353 |
| *Fusarium subglutinans* | GCF_013396075.1 | 44 | 14,039 | *Fusarium vanettenii* | GCF_000151355.1 | 51 | 15,708 |
| *Fusarium venenatum* | GCF_020744135.1 | 37 | 13,158 | *Fusarium verticillioides* | GCF_000149555.1 | 42 | 16,290 |
| *Gaeumannomyces tritici* | GCF_000145635.1 | 44 | 14,749 | *Gamsiella multidivaricata* | GCF_025024155.1 | 38 | 12,419 |
| *Gilbertella persicaria* | GCF_025201335.1 | 26 | 11,124 | *Gloeophyllum trabeum* | GCF_000344685.1 | 37 | 11,883 |
| *Halteromyces radiatus* | GCF_025201355.1 | 26 | 10,305 | *Heterobasidion irregulare* | GCF_000320585.1 | 34 | 13,270 |
| *Hyphopichia burtonii* | GCF_001661395.1 | 12 | 6,166 | *Hypoxylon fragiforme* | GCF_022984875.1 | 36 | 10,382 |
| *Hypoxylon trugodes* | GCF_022578975.1 | 39 | 12,128 | *Ilyonectria robusta* | GCF_021365365.1 | 60 | 20,715 |

Table S4: Accession ID, genome size and gene count for all species used for training the Tiberius model for **Fungi** (1/2). All annotations are NCBI RefSeq annotations.

| Species | Accession ID | Genome (Mb) | #Genes | Species | Accession ID | Genome (Mb) | #Genes |
|---|---|---|---|---|---|---|---|
| **Training (2/2)** | | | | | | | |
| *Kazachstania africana* | GCF_000304475.1 | 11 | 5,646 | *Kickxella alabastrina* | GCF_025024165.1 | 22 | 7,463 |
| *Kluyveromyces lactis* | GCF_000002515.2 | 11 | 5,335 | *Kluyveromyces marxianus* | GCF_001417885.1 | 11 | 5,151 |
| *Kockiozyma suomiensis* | GCF_038497685.1 | 14 | 6,203 | *Kockovaella imperatae* | GCF_002102565.1 | 17 | 7,428 |
| *Komagataella phaffii* | GCF_000027005.1 | 9 | 5,040 | *Kuraishia capsulata* | GCF_000576695.1 | 11 | 5,988 |
| *Kwoniella bestiolae* | GCF_000512585.2 | 25 | 9,214 | *Kwoniella botswanensis* | GCF_036426115.1 | 23 | 8,705 |
| *Kwoniella dejecticola* | GCF_000512565.2 | 24 | 8,719 | *Kwoniella europaea* | GCF_036810445.1 | 23 | 8,670 |
| *Kwoniella mangrovensis* | GCF_000507465.2 | 23 | 8,610 | *Kwoniella newhampshirensis* | GCF_039105145.1 | 20 | 7,210 |
| *Kwoniella shandongensis* | GCF_008629635.2 | 21 | 7,418 | *Kwoniella shivajii* | GCF_035658355.1 | 22 | 7,975 |
| *Lachancea thermotolerans* | GCF_000142805.1 | 10 | 5,384 | *Laetiporus sulphureus* | GCF_001632365.1 | 40 | 13,840 |
| *Lasiodiplodia theobromae* | GCF_012971845.1 | 44 | 13,054 | *Lasiosphaeria miniovina* | GCF_030549165.1 | 49 | 15,242 |
| *Lentinula edodes* | GCF_021015755.1 | 46 | 14,392 | *Limtongia smithiae* | GCF_038497785.1 | 14 | 6,123 |
| *Linderina pennispora* | GCF_002104995.1 | 26 | 9,583 | *Lindgomyces ingoldianus* | GCF_010093535.1 | 69 | 16,000 |
| *Lipomyces chichibuensis* | GCF_038497645.1 | 16 | 6,640 | *Lipomyces doorenjongii* | GCF_038497915.1 | 19 | 7,414 |
| *Lipomyces japonicus* | GCF_038497905.1 | 13 | 5,975 | *Lipomyces oligophaga* | GCF_038497815.1 | 14 | 5,861 |
| *Lipomyces tetrasporus* | GCF_029532465.1 | 21 | 8,094 | *Lobosporangium transversale* | GCF_002105155.1 | 43 | 11,983 |
| *Macroventuria anomochaeta* | GCF_010093625.1 | 33 | 13,082 | *Metarhizium brunneum* | GCF_013426205.1 | 38 | 11,555 |
| *Metarhizium robertsii* | GCF_000187425.2 | 42 | 11,688 | *Mollisia scopiformis* | GCF_001500285.1 | 49 | 18,648 |
| *Morchella importuna* | GCF_003444635.1 | 51 | 11,642 | *Mucor mucedo* | GCF_025094135.1 | 46 | 14,226 |
| *Mucor velutinosus* | GCF_035048745.1 | 40 | 13,535 | *Mycotypha africana* | GCF_025528875.1 | 29 | 11,085 |
| *Mytilinidion resinicola* | GCF_010093595.1 | 47 | 16,878 | *Myxozyma melibiosi* | GCF_037950935.1 | 15 | 6,393 |
| *Nakaseomyces glabratus* | GCF_010111755.1 | 13 | 5,508 | *Nannizzia gypsea* | GCF_000150975.2 | 23 | 9,029 |
| *Naumovozyma dairenensis* | GCF_000227115.2 | 14 | 5,814 | *Neoarthrinium moseri* | GCF_022829205.1 | 44 | 13,929 |
| *Neohortaea acidophila* | GCF_010093505.1 | 20 | 9,858 | *Neurospora crassa* | GCF_000182925.2 | 41 | 10,590 |
| *Neurospora tetraspora* | GCF_033439725.1 | 44 | 11,717 | *Paecilomyces variotii* | GCF_004022145.1 | 30 | 9,415 |
| *Paraphaeosphaeria sporulosa* | GCF_001642045.1 | 38 | 14,839 | *Parathielavia appendiculata* | GCF_033297405.1 | 33 | 12,100 |
| *Penicillium alfredii* | GCF_028826965.1 | 27 | 10,160 | *Penicillium angulare* | GCF_028827245.1 | 38 | 13,393 |
| *Penicillium argentinense* | GCF_028826775.1 | 34 | 12,141 | *Penicillium atrosanguineum* | GCF_028827265.1 | 29 | 10,995 |
| *Penicillium bovifimosum* | GCF_028826915.1 | 27 | 10,392 | *Penicillium brevicompactum* | GCF_028827555.1 | 35 | 12,371 |
| *Penicillium canariense* | GCF_028826845.1 | 32 | 10,797 | *Penicillium cataractarum* | GCF_028827025.1 | 37 | 12,826 |
| *Penicillium chrysogenum* | GCF_028827035.1 | 32 | 11,965 | *Penicillium cinerascens* | GCF_028974065.1 | 29 | 11,023 |
| *Penicillium concentricum* | GCF_028827145.1 | 30 | 11,702 | *Penicillium coprophilum* | GCF_028826855.1 | 29 | 10,975 |
| *Penicillium daleae* | GCF_028827525.1 | 40 | 12,820 | *Penicillium digitatum* | GCF_016767815.1 | 26 | 9,238 |
| *Penicillium expansum* | GCF_000769745.1 | 32 | 11,058 | *Penicillium griseofulvum* | GCF_001561935.1 | 29 | 9,628 |
| *Penicillium hispanicum* | GCF_028827665.1 | 28 | 10,111 | *Penicillium macrosclerotiorum* | GCF_028827735.1 | 34 | 11,709 |
| *Penicillium malachiteum* | GCF_028827825.1 | 36 | 13,213 | *Penicillium nucicola* | GCF_028828085.1 | 31 | 11,520 |
| *Penicillium odoratum* | GCF_028828045.1 | 32 | 12,000 | *Penicillium oxalicum* | GCF_001723175.1 | 31 | 9,966 |
| *Penicillium paradoxum* | GCF_028828445.1 | 29 | 9,841 | *Penicillium psychrosexuale* | GCF_028828465.1 | 28 | 10,478 |
| *Penicillium pulvis* | GCF_028828015.1 | 33 | 12,164 | *Penicillium robsamsonii* | GCF_028829455.1 | 29 | 11,257 |
| *Penicillium solitum* | GCF_028829755.1 | 33 | 12,723 | *Penicillium soppii* | GCF_028829465.1 | 32 | 12,167 |
| *Penicillium subrubescens* | GCF_028828155.1 | 40 | 13,459 | *Penicillium taxi* | GCF_028828555.1 | 28 | 10,052 |
| *Penicillium verhagenii* | GCF_028828195.1 | 32 | 11,667 | *Phycomyces blakesleeanus* | GCF_001638985.1 | 54 | 16,850 |
| *Phyllosticta capitalensis* | GCF_038381095.1 | 32 | 12,229 | *Phyllosticta citribraziliensis* | GCF_038025025.1 | 31 | 10,847 |
| *Plenodomus lingam* | GCF_022343315.1 | 42 | 11,989 | *Pleurotus ostreatus* | GCF_014466165.1 | 35 | 11,849 |
| *Polychytrium aggregatum* | GCF_025602895.1 | 65 | 11,051 | *Postia placenta* | GCF_002117355.1 | 42 | 12,716 |
| *Priceomyces carsonii* | GCF_900007245.1 | 12 | 6,179 | *Pseudocercospora fijiensis* | GCF_000340215.1 | 74 | 13,125 |
| *Pseudogymnoascus verrucosus* | GCF_001662655.1 | 30 | 10,799 | *Pseudomassariella vexata* | GCF_002105095.1 | 45 | 12,950 |
| *Pseudozyma flocculosa* | GCF_000417875.1 | 23 | 6,877 | *Pseudozyma hubeiensis* | GCF_000403515.1 | 18 | 7,619 |
| *Psilocybe cubensis* | GCF_017499595.1 | 46 | 13,556 | *Purpureocillium lilacinum* | GCF_023168085.1 | 39 | 10,642 |
| *Purpureocillium takamizusanense* | GCF_022605165.1 | 36 | 10,859 | *Pyricularia grisea* | GCF_004355905.1 | 45 | 12,441 |
| *Pyricularia oryzae* | GCF_000002495.2 | 41 | 13,184 | *Pyricularia pennisetigena* | GCF_004337985.1 | 49 | 11,428 |
| *Radiomyces spectabilis* | GCF_025331425.1 | 30 | 11,044 | *Ramularia collo-cygni* | GCF_900074925.1 | 32 | 11,544 |
| *Rhizophagus irregularis* | GCF_026210795.1 | 147 | 30,147 | *Rhizopus microsporus* | GCF_002708625.1 | 26 | 10,958 |
| *Rhodotorula toruloides* | GCF_000320785.1 | 20 | 8,171 | *Saccharomyces cerevisiae* | GCF_000146045.2 | 12 | 6,477 |
| *Saccharomyces kudriavzevii* | GCF_947243775.1 | 12 | 5,880 | *Saccharomyces mikatae* | GCF_947241705.1 | 12 | 5,849 |
| *Saccharomyces paradoxus* | GCF_002079055.1 | 12 | 5,840 | *Saitoella complicata* | GCF_001661265.1 | 14 | 7,200 |
| *Saprochaete ingens* | GCF_902498895.1 | 21 | 6,475 | *Scheffersomyces amazonensis* | GCF_036850825.1 | 14 | 6,209 |
| *Scheffersomyces stipitis* | GCF_000209165.1 | 15 | 5,819 | *Scheffersomyces xylosifermentans* | GCF_036884685.1 | 17 | 6,350 |
| *Schizosaccharomyces cryophilus* | GCF_000004155.1 | 12 | 5,494 | *Schizosaccharomyces japonicus* | GCF_000149845.2 | 12 | 5,224 |
| *Schizosaccharomyces octosporus* | GCF_000150505.1 | 12 | 5,347 | *Schizosaccharomyces pombe* | GCF_000002945.2 | 13 | 12,645 |
| *Sodiomyces alkalinus* | GCF_003711515.1 | 43 | 9,625 | *Sordaria macrospora* | GCF_033870435.1 | 39 | 10,804 |
| *Sparassis crispa* | GCF_003851025.1 | 39 | 13,339 | *Sphaerulina musiva* | GCF_000320565.1 | 29 | 10,228 |
| *Spizellomyces punctatus* | GCF_000182565.1 | 24 | 9,169 | *Stereum hirsutum* | GCF_000264905.1 | 47 | 14,449 |
| *Suhomyces tanzawaensis* | GCF_001661415.1 | 13 | 6,143 | *Suillus bovinus* | GCF_016758785.1 | 47 | 13,655 |
| *Suillus clintonianus* | GCF_016758775.1 | 47 | 15,647 | *Suillus fuscotomentosus* | GCF_016647785.1 | 80 | 18,660 |
| *Suillus paluster* | GCF_016628075.1 | 62 | 16,708 | *Suillus plorans* | GCF_016647745.1 | 52 | 16,525 |
| *Suillus subalutaceus* | GCF_016647625.1 | 65 | 17,233 | *Synchytrium microbalum* | GCF_006535985.1 | 26 | 6,304 |
| *Talaromyces proteolyticus* | GCF_021365285.1 | 38 | 13,665 | *Talaromyces stipitatus* | GCF_000003125.1 | 36 | 12,572 |
| *Torulaspora delbrueckii* | GCF_000243375.1 | 9 | 5,174 | *Torulaspora globosa* | GCF_014133895.1 | 9 | 5,129 |
| *Trametes versicolor* | GCF_000271585.1 | 45 | 14,562 | *Trematosphaeria pertusa* | GCF_010094035.1 | 48 | 17,364 |
| *Trichoderma aggressivum* | GCF_033847375.1 | 39 | 10,998 | *Trichoderma breve* | GCF_028502605.1 | 39 | 11,590 |
| *Trichoderma harzianum* | GCF_003025095.1 | 41 | 14,294 | *Trichoderma reesei* | GCF_000167675.1 | 33 | 9,116 |
| *Trichoderma virens* | GCF_000170995.1 | 39 | 12,382 | *Truncatella angustata* | GCF_020726525.1 | 47 | 14,971 |
| *Umbelopsis ramanniana* | GCF_025399195.1 | 23 | 10,039 | *Ustilaginoidea virens* | GCF_000687475.1 | 37 | 8,297 |
| *Ustilago hordei* | GCF_900519145.1 | 27 | 7,703 | *Ustilago maydis* | GCF_000328475.2 | 20 | 6,907 |
| *Venustampulla echinocandica* | GCF_003357145.1 | 34 | 10,707 | *Westerdykella ornata* | GCF_010094085.1 | 27 | 10,501 |
| *Wickerhamomyces anomalus* | GCF_001661255.1 | 14 | 6,571 | *Xylaria bambusicola* | GCF_022495145.1 | 46 | 12,324 |
| *Yarrowia lipolytica* | GCF_014490615.1 | 21 | 7,247 | *Zasmidium cellare* | GCF_010093935.1 | 38 | 16,072 |
| *Zychaea mexicana* | GCF_025766255.1 | 51 | 14,134 | *Zygotorulaspora mrakii* | GCF_013402915.1 | 10 | 5,246 |

Table S5: Accession ID, genome size and gene count for all species used for training the Tiberius model for **Fungi** (2/2). All annotations are NCBI RefSeq annotations.

| Species | Accession ID | Genome (Mb) | #Genes | Species | Accession ID | Genome (Mb) | #Genes |
|---|---|---|---|---|---|---|---|
| **Validation** | | | | | | | |
| *Alternaria alternata* | GCF_001642055.1 | 33 | 13,577 | *Alternaria arborescens* | GCF_004154835.1 | 34 | 12,895 |
| *Aspergillus aculeatinus* | GCF_003184765.1 | 36 | 12,350 | *Aspergillus costaricensis* | GCF_003184835.1 | 37 | 12,234 |
| *Aspergillus eucalypticola* | GCF_003184535.1 | 35 | 12,127 | *Aspergillus pseudonomiae* | GCF_009193645.1 | 38 | 13,621 |
| *Aspergillus pseudoviridinutans* | GCF_018340605.1 | 33 | 11,496 | *Aspergillus ruber* | GCF_000600275.1 | 26 | 10,229 |
| *Aureobasidium subglaciale* | GCF_000721755.1 | 26 | 10,792 | *Bipolaris sorokiniana* | GCF_000338995.1 | 34 | 12,304 |
| *Candida pseudojiufengensis* | GCF_024610245.1 | 14 | 5,846 | *Cercospora beticola* | GCF_033473495.1 | 36 | 13,212 |
| *Colletotrichum abscissum* | GCF_030869225.1 | 54 | 17,223 | *Colletotrichum chrysophilum* | GCF_026319265.1 | 56 | 14,530 |
| *Colletotrichum costaricense* | GCF_030867565.1 | 52 | 17,065 | *Colletotrichum destructivum* | GCF_034447905.1 | 52 | 15,627 |
| *Colletotrichum fructicola* | GCF_000319635.2 | 60 | 17,388 | *Coprinopsis cinerea* | GCF_000182895.1 | 36 | 13,657 |
| *Cutaneotrichosporon cavernicola* | GCF_030864355.1 | 20 | 7,753 | *Daldinia loculata* | GCF_022478755.1 | 38 | 10,988 |
| *Diaporthe citri* | GCF_014595645.1 | 64 | 15,918 | *Epithele typhae* | GCF_022376455.1 | 56 | 13,990 |
| *Fonsecaea pedrosoi* | GCF_000835455.1 | 35 | 12,573 | *Fusarium flagelliforme* | GCF_020744385.1 | 40 | 13,910 |
| *Fusarium musae* | GCF_019915245.1 | 44 | 13,962 | *Fusarium poae* | GCF_019609905.1 | 44 | 14,125 |
| *Geosmithia morbida* | GCF_012550715.1 | 27 | 7,826 | *Glarea lozoyensis* | GCF_000409485.1 | 40 | 13,081 |
| *Guyanagaster necrorhizus* | GCF_019112545.1 | 54 | 14,487 | *Kalmanozyma brasiliensis* | GCF_000497045.1 | 17 | 6,542 |
| *Kwoniella dendrophila* | GCF_036810415.1 | 23 | 8,131 | *Kwoniella pini* | GCF_000512605.1 | 21 | 7,931 |
| *Lichtheimia ornata* | GCF_029851405.1 | 38 | 13,169 | *Lipomyces arxii* | GCF_038497795.1 | 12 | 5,792 |
| *Marasmius oreades* | GCF_018924745.1 | 44 | 13,890 | *Microdochium trichocladiopsis* | GCF_020744255.1 | 49 | 16,222 |
| *Moesziomyces antarcticus* | GCF_000747765.1 | 18 | 6,885 | *Morchella sextelata* | GCF_020137385.1 | 54 | 13,182 |
| *Naumovozyma castellii* | GCF_000237345.1 | 11 | 5,867 | *Neurospora hispaniola* | GCF_033458365.1 | 41 | 10,876 |
| *Paracoccidioides lutzii* | GCF_000150705.2 | 33 | 8,953 | *Penicilliopsis zonata* | GCF_001890105.1 | 26 | 10,027 |
| *Penicillium antarcticum* | GCF_028974205.1 | 31 | 11,217 | *Penicillium crustosum* | GCF_028827405.1 | 33 | 12,327 |
| *Penicillium diatomitis* | GCF_028827545.1 | 34 | 9,582 | *Penicillium hordei* | GCF_028827395.1 | 34 | 12,337 |
| *Penicillium longicatenatum* | GCF_028827895.1 | 32 | 11,974 | *Penicillium rubens* | GCF_028828025.1 | 30 | 11,613 |
| *Penicillium verrucosum* | GCF_028828655.1 | 33 | 11,554 | *Penicillium vulpinum* | GCF_028829585.1 | 32 | 11,510 |
| *Pisolithus orientalis* | GCF_024613125.1 | 58 | 12,436 | *Rhinocladiella mackenziei* | GCF_000835555.1 | 32 | 11,418 |
| *Rhodofomes roseus* | GCF_022264815.1 | 38 | 13,143 | *Schizophyllum commune* | GCF_000143185.2 | 39 | 16,467 |
| *Schizosaccharomyces osmophilus* | GCF_027921745.1 | 11 | 5,574 | *Suillus discolor* | GCF_016758755.1 | 61 | 16,641 |
| *Suillus subaureus* | GCF_016647635.1 | 58 | 15,900 | *Tetrapisispora phaffii* | GCF_000236905.1 | 12 | 5,462 |
| *Wickerhamomyces ciferrii* | GCF_000313485.1 | 16 | 6,702 | *Zygosaccharomyces rouxii* | GCF_000026365.1 | 10 | 5,323 |
| **Testing** | | | | | | | |
| *Agaricus bisporus* | GCF_000300555.1 | 33 | 11,443 | *Cryphonectria parasitica* | GCF_011745365.1 | 44 | 11,709 |
| *Parastagonospora nodorum* | GCF_000146915.1 | 37 | 16,114 | *Puccinia striiformis* | GCF_021901695.1 | 89 | 17,813 |
| *Punctularia strigosozonata* | GCF_000264995.1 | 34 | 11,644 | *Tilletiopsis washingtonensis* | GCF_003144115.1 | 19 | 7,053 |

Table S6: Accession ID, genome size and gene count for all species used for validating and testing the Tiberius model for **Fungi**. All annotations are NCBI RefSeq annotations.

| Species | Accession ID | Genome (Mb) | Annotation | #Genes | Species | Accession ID | Genome (Mb) | Annotation | #Genes |
|---|---|---|---|---|---|---|---|---|---|
| **Training** | | | | | | | | | |
| *Aedes aegypti* | GCF_002204515.2 | 1,279 | RefSeq | 14,731 | *Agrotis ipsilon* | GCA_028554685.2 | 515 | BRAKER | 11,962 |
| *Amphipyra tragopoginis* | GCA_905220435.1 | 806 | BRAKER | 18,183 | *Anabrus simplex* | GCF_040414725.1 | 6,424 | RefSeq | 14,968 |
| *Anastrepha ludens* | GCF_028408465.1 | 821 | BRAKER | 18,103 | *Andrena minutula* | GCA_929113495.1 | 380 | BRAKER | 12,657 |
| *Anopheles arabiensis* | GCF_016920715.1 | 257 | BRAKER | 12,969 | *Anopheles farauti* | GCA_000473445.2 | 183 | BRAKER | 11,884 |
| *Anopheles stephensi* | GCF_013141755.1 | 243 | BRAKER | 12,990 | *Aphis glycines* | GCA_009761285.1 | 308 | BRAKER | 16,004 |
| *Apis mellifera* | GCF_003254395.2 | 225 | BRAKER | 11,040 | *Aporophyla lueneburgensis* | GCA_932294355.1 | 978 | BRAKER | 23,312 |
| *Arctia plantaginis* | GCA_902825455.1 | 578 | BRAKER | 16,231 | *Armigeres subalbatus* | GCF_024139115.2 | 1,336 | RefSeq | 19,126 |
| *Bactrocera dorsalis* | GCF_023373825.1 | 530 | BRAKER | 18,592 | *Bactrocera minax* | GCA_021498325.1 | 324 | BRAKER | 12,187 |
| *Bactrocera oleae* | GCF_001188975.3 | 485 | BRAKER | 14,673 | *Bactrocera tryoni* | GCF_016617805.1 | 571 | BRAKER | 17,801 |
| *Bemisia tabaci* | GCF_001854935.1 | 615 | BRAKER | 14,377 | *Bicyclus anynana* | GCF_947172395.1 | 457 | RefSeq | 13,653 |
| *Bombus impatiens* | GCF_000188095.3 | 247 | BRAKER | 12,389 | *Bradysia coprophila* | GCA_014529535.2 | 310 | BRAKER | 19,444 |
| *Calliphora vicina* | GCA_958450345.1 | 707 | BRAKER | 16,315 | *Camponotus floridanus* | GCF_003227725.1 | 284 | BRAKER | 13,038 |
| *Cardiocondyla obscurior* | GCA_019399895.1 | 193 | BRAKER | 15,059 | *Ceratitis capitata* | GCF_000347755.3 | 436 | BRAKER | 13,455 |
| *Chilo suppressalis* | GCA_902850365.2 | 783 | BRAKER | 22,590 | *Chrysoperla carnea* | GCF_905475395.1 | 560 | BRAKER | 14,441 |
| *Cimex lectularius* | GCF_000648675.2 | 511 | BRAKER | 13,021 | *Cochliomyia hominivorax* | GCA_004302925.2 | 534 | BRAKER | 12,788 |
| *Cryptotermes secundus* | GCF_002891405.2 | 1,019 | BRAKER | 14,614 | *Culex quinquefasciatus* | GCF_015732765.1 | 573 | BRAKER | 15,246 |
| *Dendroctonus ponderosae* | GCF_020466585.1 | 224 | BRAKER | 14,204 | *Dermacentor andersoni* | GCF_023375885.2 | 2,425 | RefSeq | 20,700 |
| *Diuraphis noxia* | GCF_001186385.1 | 395 | BRAKER | 23,559 | *Drosophila birchii* | GCA_008042755.1 | 157 | BRAKER | 16,937 |
| *Drosophila grimshawi* | GCF_018153295.1 | 191 | BRAKER | 12,744 | *Drosophila innubila* | GCA_004354385.2 | 166 | BRAKER | 15,008 |
| *Drosophila navojoa* | GCF_001654015.2 | 147 | BRAKER | 12,770 | *Drosophila obscura* | GCF_018151105.1 | 180 | BRAKER | 16,091 |
| *Drosophila suzukii* | GCF_013340165.1 | 268 | BRAKER | 17,364 | *Ennomos fuscantarius* | GCA_905220475.3 | 445 | BRAKER | 14,282 |
| *Episyrphus balteatus* | GCF_945859705.1 | 535 | BRAKER | 14,004 | *Formica exsecta* | GCF_003651465.1 | 278 | BRAKER | 18,914 |
| *Frankliniella occidentalis* | GCF_000697945.3 | 406 | BRAKER | 22,507 | *Galleria mellonella* | GCF_026898425.1 | 472 | BRAKER | 13,576 |
| *Glossina palpalis* | GCA_000818775.1 | 380 | BRAKER | 9,740 | *Gryllus bimaculatus* | GCA_017312745.1 | 1,658 | BRAKER | 11,991 |
| *Halyomorpha halys* | GCA_000696795.3 | 1,150 | BRAKER | 16,536 | *Harpegnathos saltator* | GCF_003227715.2 | 334 | BRAKER | 14,955 |
| *Helicoverpa zea* | GCF_022581195.2 | 375 | BRAKER | 16,321 | *Hermetia illucens* | GCF_905115235.1 | 1,005 | BRAKER | 15,393 |
| *Hormaphis cornu* | GCA_017140985.1 | 320 | BRAKER | 14,080 | *Hycleus cichorii* | GCA_013841215.1 | 99 | BRAKER | 8,417 |
| *Ips nitidus* | GCA_018691245.2 | 231 | BRAKER | 21,165 | *Laodelphax striatellus* | GCA_017141395.1 | 510 | BRAKER | 13,701 |
| *Lasiommata megera* | GCA_928268935.1 | 488 | BRAKER | 12,772 | *Leptopilina boulardi* | GCF_019393585.1 | 355 | BRAKER | 14,452 |
| *Lymantria dispar* | GCA_016802235.1 | 998 | BRAKER | 19,022 | *Malaya genurostris* | GCF_030247185.1 | 831 | RefSeq | 13,645 |
| *Megalopta genalis* | GCF_011865705.1 | 390 | BRAKER | 10,806 | *Melanaphis sacchari* | GCF_002803265.2 | 300 | BRAKER | 13,106 |
| *Melitaea cinxia* | GCF_905220565.1 | 499 | BRAKER | 19,514 | *Metopolophium dirhodum* | GCA_019925205.1 | 481 | BRAKER | 22,550 |
| *Monomorium pharaonis* | GCF_013373865.1 | 326 | BRAKER | 17,086 | *Musca domestica* | GCF_000371365.1 | 750 | BRAKER | 13,580 |
| *Mythimna separata* | GCA_029852925.1 | 688 | BRAKER | 17,248 | *Nasonia vitripennis* | GCF_009193385.2 | 297 | BRAKER | 21,342 |
| *Nezara viridula* | GCA_928085145.1 | 1,185 | BRAKER | 19,624 | *Nicrophorus vespilloides* | GCF_001412225.1 | 195 | BRAKER | 13,883 |
| *Onthophagus taurus* | GCF_000648695.1 | 267 | BRAKER | 19,160 | *Ooceraea biroi* | GCF_003672135.1 | 224 | BRAKER | 14,589 |
| *Operophtera brumata* | GCA_932527175.1 | 619 | BRAKER | 15,652 | *Oryctes rhinoceros* | GCA_020654165.1 | 377 | BRAKER | 11,931 |
| *Ostrinia furnacalis* | GCF_004193835.2 | 435 | BRAKER | 15,189 | *Pediculus humanus* | GCF_000006295.1 | 111 | BRAKER | 8,810 |
| *Periplaneta americana* | GCA_025594305.2 | 3,056 | BRAKER | 19,261 | *Phlebotomus papatasi* | GCF_024763615.1 | 352 | BRAKER | 10,881 |
| *Plodia interpunctella* | GCF_027563975.1 | 291 | BRAKER | 14,816 | *Pogonomyrmex californicus* | GCA_024349325.1 | 252 | BRAKER | 28,178 |
| *Polypedilum vanderplanki* | GCA_018290095.1 | 119 | BRAKER | 14,780 | *Propsilocerus akamusi* | GCA_018397935.1 | 86 | BRAKER | 12,517 |
| *Rhogogaster chlorosoma* | GCA_944452935.1 | 255 | BRAKER | 21,737 | *Rhopalosiphum padi* | GCA_020882245.1 | 338 | BRAKER | 13,277 |
| *Rhynchophorus ferrugineus* | GCA_030347505.1 | 779 | BRAKER | 12,216 | *Scaptomyza flava* | GCA_030179655.1 | 330 | BRAKER | 14,660 |
| *Schizaphis graminum* | GCA_020882235.1 | 499 | BRAKER | 28,035 | *Schlechtendalia chinensis* | GCA_019022885.1 | 280 | BRAKER | 17,424 |
| *Sipha flava* | GCF_003268045.1 | 353 | BRAKER | 9,105 | *Sitobion avenae* | GCA_019425605.1 | 393 | BRAKER | 10,662 |
| *Sitophilus oryzae* | GCF_002938485.1 | 771 | BRAKER | 20,600 | *Sogatella furcifera* | GCA_017141385.1 | 564 | BRAKER | 11,007 |
| *Solenopsis invicta* | GCF_016802725.1 | 378 | BRAKER | 18,058 | *Spodoptera litura* | GCF_002706865.2 | 430 | BRAKER | 16,178 |
| *Stomoxys calcitrans* | GCF_963082655.1 | 1,071 | RefSeq | 15,234 | *Temnothorax longispinosus* | GCA_030848805.1 | 298 | BRAKER | 20,311 |
| *Thrips palmi* | GCF_012932325.1 | 238 | BRAKER | 14,785 | *Thymelicus sylvestris* | GCA_911387775.1 | 471 | BRAKER | 15,050 |
| *Trichoplusia ni* | GCF_003590095.1 | 368 | BRAKER | 15,313 | *Zeugodacus cucurbitae* | GCF_028554725.1 | 439 | BRAKER | 15,987 |
| **Test** | | | | | | | | | |
| *Bombyx mori* | GCF_030269925.1 | 462 | RefSeq | 18,210 | *Cataglyphis hispanica* | GCF_021464435.1 | 206 | RefSeq | 11,502 |
| *Colias croceus* | GCF_905220415.1 | 325 | RefSeq | 15,772 | *Danaus plexippus* | GCF_018135715.1 | 245 | RefSeq | 14,790 |
| *Drosophila melanogaster* | GCF_000001215.4 | 144 | RefSeq | 17,872 | *Leptidea sinapis* | GCF_905404315.1 | 686 | RefSeq | 15,767 |
| *Nymphalis io* | GCF_905147045.1 | 384 | RefSeq | 13,823 | *Osmia bicornis* | GCF_907164935.1 | 223 | RefSeq | 12,706 |
| *Tribolium castaneum* | GCF_031307605.1 | 242 | RefSeq | 15,518 | *Vanessa cardui* | GCF_905220365.1 | 425 | RefSeq | 15,000 |
| *Zerene cesonia* | GCF_012273895.1 | 266 | RefSeq | 12,772 | | | | | |
| **Validation** | | | | | | | | | |
| *Anopheles marshallii* | GCF_943734725.1 | 226 | RefSeq | 13,358 | *Drosophila ananassae* | GCF_017639315.1 | 214 | BRAKER | 16,982 |
| *Drosophila virilis* | GCF_030788295.1 | 189 | BRAKER | 15,650 | *Tenebrio molitor* | GCF_963966145.1 | 277 | BRAKER | 19,578 |

Table S7: Accession ID, genome size and gene count for all species used for training, validating and testing the Tiberius model for **Insecta**. Annotation source refers to the annotation used (BRAKER = AUGUSTUS/GeneMark pipeline; or RefSeq).

| Species | Accession ID | Genome (Mb) | #Genes | Species | Accession ID | Genome (Mb) | #Genes |
|---|---|---|---|---|---|---|---|
| **Training** | | | | | | | |
| *Acipenser ruthenus* | GCF_902713425.1 | 1,900 | 80,041 | *Alligator mississippiensis* | GCF_030867095.1 | 2,347 | 31,016 |
| *Alosa sapidissima* | GCF_018492685.1 | 904 | 47,628 | *Amblyraja radiata* | GCF_010909765.2 | 2,559 | 25,978 |
| *Anguilla anguilla* | GCF_013347855.1 | 979 | 31,451 | *Anolis sagrei* | GCF_037176765.1 | 1,951 | 23,363 |
| *Apus apus* | GCF_020740795.1 | 1,100 | 16,724 | *Bombina bombina* | GCF_027579735.1 | 10,020 | 31,613 |
| *Carcharodon carcharias* | GCF_017639515.1 | 4,286 | 25,164 | *Corvus hawaiiensis* | GCF_020740725.1 | 1,152 | 21,647 |
| *Dendropsophus ebraccatus* | GCF_027789765.1 | 2,215 | 37,424 | *Denticeps clupeoides* | GCF_900700375.1 | 567 | 40,127 |
| *Elgaria multicarinata webbii* | GCF_023053635.1 | 1,791 | 21,748 | *Emys orbicularis* | GCF_028017835.1 | 2,310 | 22,788 |
| *Erpetoichthys calabaricus* | GCF_900747795.2 | 3,614 | 27,285 | *Esox lucius* | GCF_011004845.1 | 919 | 30,105 |
| *Eublepharis macularius* | GCF_028583425.1 | 2,237 | 24,025 | *Falco peregrinus* | GCF_023634155.1 | 1,312 | 19,918 |
| *Gavia stellata* | GCF_030936135.1 | 1,321 | 17,980 | *Labrus mixtus* | GCF_963584025.1 | 741 | 41,941 |
| *Latimeria chalumnae* | GCF_037176945.1 | 2,948 | 28,938 | *Lepisosteus oculatus* | GCF_040954835.1 | 1,201 | 49,180 |
| *Megalops cyprinoides* | GCF_013368585.1 | 959 | 27,986 | *Mobula birostris* | GCF_030028105.1 | 4,015 | 27,860 |
| *Narcine bancroftii* | GCF_036971445.1 | 3,328 | 30,148 | *Osmerus eperlanus* | GCF_963692335.1 | 509 | 34,446 |
| *Parambassis ranga* | GCF_900634625.1 | 551 | 27,951 | *Pelobates fuscus* | GCF_036172605.1 | 3,606 | 49,915 |
| *Podarcis raffonei* | GCF_027172205.1 | 1,513 | 26,204 | *Rana temporaria* | GCF_905171775.1 | 4,111 | 38,300 |
| *Rhinatrema bivittatum* | GCF_901001135.1 | 5,319 | 28,114 | *Scleropages formosus* | GCF_900964775.1 | 785 | 27,406 |
| *Solea solea* | GCF_958295425.1 | 644 | 37,145 | *Sparus aurata* | GCF_900880675.1 | 834 | 32,006 |
| *Aotus nancymaae* | GCF_000952055.2 | 2,862 | 31,334 | *Camelus bactrianus* | GCF_000767855.1 | 1,993 | 29,764 |
| *Canis lupus familiaris* | GCF_000002285.3 | 2,411 | 36,943 | *Cavia porcellus* | GCF_000151735.1 | 2,723 | 31,557 |
| *Cebus capucinus* | GCF_001604975.1 | 2,718 | 36,525 | *Ceratotherium simum* | GCF_000283155.1 | 2,464 | 25,301 |
| *Chinchilla lanigera* | GCF_000276665.1 | 2,391 | 30,122 | *Condylura cristata* | GCF_000260355.1 | 1,770 | 19,773 |
| *Desmodus rotundus* | GCF_002940915.1 | 2,064 | 29,514 | *Dipodomys ordii* | GCF_000151885.1 | 2,236 | 23,237 |
| *Enhydra lutris* | GCF_002288905.1 | 2,455 | 24,129 | *Eptesicus fuscus* | GCF_000308155.1 | 2,027 | 30,638 |
| *Equus caballus* | GCF_000002305.2 | 2,475 | 26,780 | *Heterocephalus glaber* | GCF_000247695.1 | 2,618 | 37,544 |
| *Ictidomys tridecemlineatus* | GCF_000236235.1 | 2,478 | 26,378 | *Jaculus jaculus* | GCF_000280705.1 | 2,835 | 25,112 |
| *Loxodonta africana* | GCF_000001905.1 | 3,197 | 28,964 | *Marmota marmota* | GCF_001458135.1 | 2,511 | 27,672 |
| *Microcebus murinus* | GCF_000165445.2 | 2,487 | 31,964 | *Microtus ochrogaster* | GCF_000317375.1 | 2,287 | 26,397 |
| *Mus musculus* | GCF_000001635.26 | 2,731 | 49,997 | *Mus pahari* | GCF_900095145.1 | 2,475 | 29,483 |
| *Neomonachus schauinslandi* | GCF_002201575.1 | 2,401 | 24,767 | *Ochotona princeps* | GCF_000292845.1 | 2,230 | 21,216 |
| *Octodon degus* | GCF_000260255.1 | 2,996 | 39,965 | *Odobenus rosmarus* | GCF_000321225.1 | 2,400 | 24,996 |
| *Otolemur garnettii* | GCF_000181295.1 | 2,520 | 31,917 | *Panthera pardus* | GCF_001857705.1 | 2,578 | 34,200 |
| *Propithecus coquereli* | GCF_000956105.1 | 2,798 | 22,796 | *Puma concolor* | GCF_003327715.1 | 2,433 | 23,982 |
| *Rattus norvegicus* | GCF_000001895.5 | 2,870 | 39,469 | *Saimiri boliviensis* | GCF_000235385.1 | 2,609 | 27,148 |
| *Sorex araneus* | GCF_000181275.1 | 2,423 | 21,793 | *Trichechus manatus* | GCF_000243295.1 | 3,104 | 25,210 |
| **Testing** | | | | | | | |
| *Archocentrus centrarchus* | GCF_007364275.1 | 933 | 29,275 | *Betta splendens* | GCF_900634795.4 | 427 | 28,938 |
| *Gallus gallus* | GCF_016700215.2 | 1,051 | 25,564 | *Pristiophorus japonicus* | GCF_044704955.1 | 5,923 | 55,857 |
| *Takifugu rubripes* | GCF_901000725.2 | 384 | 27,434 | *Zootoca vivipara* | GCF_963506605.1 | 1,440 | 22,823 |

Table S8: Accession ID, genome size and gene count for all species according to the corresponding NCBI RefSeq annotation used for training and testing the Tiberius model for **Vertebrata**.

| Species | Accession ID | Genome (Mb) | #Genes |
|---|---|---|---|
| **Testing** | | | |
| *Bos taurus* | GCF_002263795.3 | 2,771 | 37,063 |
| *Delphinapterus leucas* | GCF_002288925.2 | 2,363 | 27,561 |
| *Homo sapiens* | GCF_000001405.40 | 3,099 | 59,792 |

Table S9: Accession ID, genome size and gene count for all species according to the corresponding NCBI RefSeq annotation used for testing the Tiberius model for **Mammalia**. The Mammalia training and validation is described in the original Tiberius paper [6] and remained unchanged.

Table S10: Runtime benchmark for Chlorophyta.

| Species | Tool | Runtime (h:mm) |
|---|---|---|
| *Bathycoccus prasinos* | Helixer | 0:04 |
| | Tiberius | 0:01 |
| | BRAKER3 | 10:34 |
| *Edaphochlamys debaryana* | Helixer | 0:25 |
| | Tiberius | 0:04 |
| | BRAKER3 | 24:24 |

Table S11: Runtime benchmark for Bacillariophyta.

| Species | Tool | Runtime (h:mm) |
|---|---|---|
| *Phaeodactylum tricornutum* | Tiberius | 0:01 |
| | BRAKER3 | 19:57 |
| *Thalassiosira pseudonana* | Tiberius | 0:01 |
| | BRAKER3 | 18:30 |

Table S12: Runtime benchmark for Mesangiospermae.

| Species | Tool | Runtime (h:mm) |
|---|---|---|
| *Arabidopsis thaliana* | Helixer | 0:15 |
| | ANNEVO | 0:05 |
| | Tiberius | 0:04 |
| | BRAKER3 | 33:02 |
| *Brachypodium distachyon* | Helixer | 0:38 |
| | ANNEVO | 0:10 |
| | Tiberius | 0:07 |
| | BRAKER3 | 4:23 |
| *Eschscholzia californica* | Helixer | 0:53 |
| | ANNEVO | 0:12 |
| | Tiberius | 0:10 |
| | BRAKER3 | 31:29 |
| *Medicago truncatula* | Helixer | 0:59 |
| | ANNEVO | 0:14 |
| | Tiberius | 0:11 |
| | BRAKER3 | 5:24 |
| *Freycinetia multiflora* | Helixer | 1:23 |
| | ANNEVO | 0:12 |
| | Tiberius | 0:13 |
| | BRAKER3 | 6:02 |
| *Mimulus guttatus* | Helixer | 0:41 |
| | ANNEVO | 0:09 |
| | Tiberius | 0:08 |
| | BRAKER3 | 46:36 |
| *Urochloa brizantha* | Helixer | 6:16 |
| | ANNEVO | 1:34 |
| | Tiberius | 1:04 |
| | BRAKER3 | 16:06 |

Table S13: Runtime benchmark for Insecta.

| Species | Tool | Runtime (h:mm) |
|---|---|---|
| *Bombyx mori* | Helixer | 1:39 |
| | ANNEVO | 0:15 |
| | Tiberius | 0:16 |
| | BRAKER3 | 63:28 |
| *Cataglyphis hispanica* | Helixer | 0:52 |
| | ANNEVO | 0:07 |
| | Tiberius | 0:08 |
| | BRAKER3 | 36:48 |
| *Colias croceus* | Helixer | 1:12 |
| | ANNEVO | 0:10 |
| | Tiberius | 0:11 |
| | BRAKER3 | 51:57 |
| *Danaus plexippus* | Helixer | 0:49 |
| | ANNEVO | 0:09 |
| | Tiberius | 0:09 |
| | BRAKER3 | 46:13 |
| *Drosophila melanogaster* | Helixer | 0:54 |
| | ANNEVO | 0:07 |
| | Tiberius | 0:09 |
| | BRAKER3 | 34:32 |
| *Leptidea sinapis* | Helixer | 2:31 |
| | ANNEVO | 0:21 |
| | Tiberius | 0:22 |
| | BRAKER3 | 70:55 |
| *Nymphalis io* | Helixer | 1:20 |
| | ANNEVO | 0:13 |
| | Tiberius | 0:13 |
| | BRAKER3 | 55:11 |
| *Osmia bicornis* | Helixer | 0:52 |
| | ANNEVO | 0:08 |
| | Tiberius | 0:10 |
| | BRAKER3 | 25:45 |
| *Tribolium castaneum* | Helixer | 0:57 |
| | ANNEVO | 0:09 |
| | Tiberius | 0:09 |
| | BRAKER3 | 40:03 |
| *Vanessa cardui* | Helixer | 1:29 |
| | ANNEVO | 0:13 |
| | Tiberius | 0:15 |
| | BRAKER3 | 50:12 |
| *Zerene cesonia* | Helixer | 1:01 |
| | ANNEVO | 0:10 |
| | Tiberius | 0:10 |
| | BRAKER3 | 27:41 |

Table S14: Runtime benchmark for Fungi.

| Species | Tool | Runtime (h:mm) |
|---|---|---|
| *Agaricus bisporus* | Helixer | 0:08 |
| | ANNEVO | 0:03 |
| | Tiberius | 0:02 |
| | BRAKER3 | 1:38 |
| *Cryphonectria parasitica* | Helixer | 0:07 |
| | ANNEVO | 0:02 |
| | Tiberius | 0:01 |
| | BRAKER3 | 1:10 |
| *Parastagonospora nodorum* | Helixer | 0:06 |
| | ANNEVO | 0:02 |
| | Tiberius | 0:01 |
| | BRAKER3 | 1:30 |
| *Puccinia striiformis* | Helixer | 0:15 |
| | ANNEVO | 0:02 |
| | Tiberius | 0:02 |
| | BRAKER3 | 2:11 |
| *Punctularia strigosozonata* | Helixer | 0:06 |
| | ANNEVO | 0:02 |
| | Tiberius | 0:01 |
| | BRAKER3 | 1:55 |
| *Tilletiopsis washingtonensis* | Helixer | 0:03 |
| | ANNEVO | 0:01 |
| | Tiberius | 0:01 |
| | BRAKER3 | 1:41 |

Table S15: Runtime benchmark for Mammalia.

| Species | Tool | Runtime (h:mm) |
|---|---|---|
| *Bos taurus* | Helixer | 10:53 |
| | ANNEVO | 1:24 |
| | Tiberius | 1:20 |
| | BRAKER3 | 83:21 |
| *Delphinapterus leucas* | Helixer | 11:10 |
| | ANNEVO | 1:18 |
| | Tiberius | 1:16 |
| | BRAKER3 | 76:40 |
| *Homo sapiens* | Helixer | 10:56 |
| | ANNEVO | 1:55 |
| | Tiberius | 1:27 |
| | BRAKER3 | 71:22 |

Table S16: Runtime benchmark for Vertebrata.

| Species | Tool | Runtime (h:mm) |
|---|---|---|
| *Archocentrus centrarchus* | Helixer | 3:43 |
| | ANNEVO | 0:38 |
| | Tiberius | 0:36 |
| | BRAKER3 | 40:26 |
| *Betta splendens* | Helixer | 1:39 |
| | ANNEVO | 0:17 |
| | Tiberius | 0:18 |
| | BRAKER3 | 46:52 |
| *Gallus gallus* | Helixer | 4:21 |
| | ANNEVO | 0:44 |
| | Tiberius | 0:39 |
| | BRAKER3 | 64:34 |
| *Pristiophorus japonicus* | Helixer | 21:24 |
| | ANNEVO | 4:16 |
| | Tiberius | 3:12 |
| | BRAKER3 | 34:11 |
| *Takifugu rubripes* | Helixer | 1:29 |
| | ANNEVO | 0:12 |
| | Tiberius | 0:17 |
| | BRAKER3 | 53:25 |
| *Zootoca vivipara* | Helixer | 6:06 |
| | ANNEVO | 0:52 |
| | Tiberius | 0:49 |
| | BRAKER3 | 64:03 |

| Clade | Species | Helixer | ANNEVO |
|---|---|---|---|
| Mesangiospermae | *Arabidopsis thaliana* | train+val | test |
| Mesangiospermae | *Brachypodium distachyon* | – | – |
| Mesangiospermae | *Eschscholzia californica* | – | – |
| Mesangiospermae | *Freycinetia multiflora* | – | – |
| Mesangiospermae | *Mimulus guttatus* | train+val | – |
| Mesangiospermae | *Medicago truncatula* | – | – |
| Mesangiospermae | *Urochloa brizantha* | – | – |
| Fungi | *Agaricus bisporus* | train+val | test |
| Fungi | *Cryphonectria parasitica* | test | test |
| Fungi | *Parastagonospora nodorum* | train+val | test |
| Fungi | *Puccinia striiformis* | – | test |
| Fungi | *Punctularia strigosozonata* | train+val | test |
| Fungi | *Tilletiopsis washingtonensis* | train+val | test |
| Insecta | *Bombyx mori* | train+val | test |
| Insecta | *Cataglyphis hispanica* | – | train |
| Insecta | *Colias croceus* | test | test |
| Insecta | *Danaus plexippus* | test | – |
| Insecta | *Drosophila melanogaster* | train+val | test |
| Insecta | *Leptidea sinapis* | – | test |
| Insecta | *Nymphalis io* | – | – |
| Insecta | *Osmia bicornis* | train+val | – |
| Insecta | *Tribolium castaneum* | test | – |
| Insecta | *Vanessa cardui* | – | train |
| Insecta | *Zerene cesonia* | train+val | – |
| Mammalia | *Bos taurus* | test | test |
| Mammalia | *Delphinapterus leucas* | train+val | – |
| Mammalia | *Homo sapiens* | train+val | test |
| Vertebrata | *Archocentrus centrarchus* | train+val | test |
| Vertebrata | *Betta splendens* | train+val | test |
| Vertebrata | *Gallus gallus* | train+val | test |
| Vertebrata | *Pristiophorus japonicus* | train+val | – |
| Vertebrata | *Takifugu rubripes* | train+val | test |
| Vertebrata | *Zootoca vivipara* | test | test |

Table S17: Usage of Tiberius test species during the training of the Helixer and ANNEVO models. None of the species were used to validate or train Tiberius.

Table S18: Average accuracy benchmark for all test species of each Tiberius model, compared to *ab initio* prediction of Helixer and ANNEVO, and evidence supported prediction of BRAKER3, where BRAKER3 used the proteomes of Tiberius' training species and short-read RNA-Seq.

| Clade | Tool | $n$ | Exon | | | Transcript | | | Gene | | |
|---|---|---|---|---|---|---|---|---|---|---|---|
| | | | Sn | Pr | F1 | Sn | Pr | F1 | Sn | Pr | F1 |
| Chlorophyta | Helixer | 2 | 69.5 | 64.2 | 66.7 | 43.2 | 43.4 | 43.0 | 43.2 | 43.4 | 43.0 |
| | Tiberius | 2 | 66.3 | 67.0 | 66.6 | 44.7 | 44.9 | 44.7 | 44.7 | 44.9 | 44.7 |
| | BRAKER3 | 2 | 51.2 | 66.8 | 57.6 | 42.0 | 46.8 | 44.3 | 42.1 | 55.7 | 47.9 |
| Bacillariophyta | Tiberius | 2 | 66.8 | 75.0 | 70.6 | 60.2 | 66.2 | 63.1 | 62.4 | 66.2 | 64.2 |
| | BRAKER3 | 2 | 60.0 | 72.5 | 65.6 | 57.1 | 60.2 | 58.5 | 59.0 | 72.9 | 65.1 |
| Mesangiospermae | Helixer | 7 | 84.7 | 75.8 | 79.7 | 50.7 | 54.6 | 52.1 | 64.6 | 54.6 | 59.0 |
| | ANNEVO | 7 | 80.8 | 83.7 | 82.2 | 45.8 | 60.7 | 52.0 | 58.4 | 60.7 | 59.5 |
| | Tiberius | 7 | 83.9 | 89.5 | 86.6 | 56.7 | 69.9 | 62.3 | 72.5 | 69.9 | 71.0 |
| | BRAKER3 | 7 | 80.6 | 92.4 | 86.0 | 59.2 | 73.3 | 65.2 | 73.7 | 79.0 | 76.0 |
| Fungi | Helixer | 6 | 64.9 | 53.8 | 58.7 | 32.2 | 26.6 | 29.0 | 32.2 | 26.6 | 29.0 |
| | ANNEVO | 6 | 66.1 | 62.8 | 64.3 | 35.6 | 35.3 | 35.4 | 35.6 | 35.3 | 35.4 |
| | Tiberius | 6 | 70.9 | 70.9 | 70.9 | 44.7 | 46.1 | 45.3 | 44.7 | 46.1 | 45.3 |
| | BRAKER3 | 6 | 63.8 | 74.5 | 68.6 | 43.8 | 48.5 | 45.8 | 43.8 | 54.7 | 48.5 |
| Insecta | Helixer | 11 | 79.5 | 72.8 | 75.9 | 27.8 | 36.9 | 31.6 | 42.5 | 36.9 | 39.5 |
| | ANNEVO | 11 | 80.5 | 80.6 | 80.5 | 31.8 | 47.3 | 38.0 | 48.7 | 47.3 | 47.9 |
| | Tiberius | 11 | 83.2 | 85.7 | 84.4 | 44.6 | 50.5 | 47.1 | 68.1 | 50.5 | 57.6 |
| | BRAKER3 | 11 | 82.1 | 92.1 | 86.7 | 56.8 | 67.1 | 61.3 | 76.4 | 74.7 | 75.4 |
| Mammalia | Helixer | 3 | 77.4 | 71.0 | 74.1 | 10.4 | 24.0 | 14.5 | 24.6 | 24.0 | 24.3 |
| | ANNEVO | 3 | 81.5 | 80.3 | 80.9 | 18.8 | 36.6 | 24.8 | 44.4 | 36.6 | 40.1 |
| | Tiberius | 3 | 83.4 | 90.7 | 86.9 | 27.3 | 59.5 | 37.4 | 64.6 | 59.5 | 61.9 |
| | BRAKER3 | 3 | 69.3 | 93.7 | 79.6 | 33.9 | 66.2 | 44.8 | 68.9 | 72.6 | 70.4 |
| Vertebrata | Helixer | 6 | 78.7 | 68.2 | 72.9 | 12.8 | 21.0 | 15.7 | 23.3 | 21.0 | 22.1 |
| | ANNEVO | 6 | 81.6 | 80.0 | 80.7 | 21.7 | 36.6 | 26.8 | 38.9 | 36.6 | 37.6 |
| | Tiberius | 6 | 85.6 | 86.2 | 85.8 | 33.6 | 51.9 | 39.8 | 60.4 | 51.9 | 55.3 |
| | BRAKER3 | 6 | 76.8 | 95.2 | 84.9 | 45.1 | 70.1 | 54.5 | 69.5 | 80.1 | 74.3 |
| Overall | Helixer | 33 | 77.6 | 69.0 | 72.9 | 29.1 | 34.7 | 31.0 | 40.2 | 34.7 | 37.2 |
| | ANNEVO | 33 | 78.2 | 77.9 | 78.0 | 32.4 | 45.1 | 37.2 | 46.2 | 45.1 | 45.5 |
| | Tiberius | 33 | 81.6 | 84.4 | 82.9 | 43.6 | 54.9 | 47.8 | 63.1 | 54.9 | 58.2 |
| | BRAKER3 | 33 | 76.3 | 89.7 | 82.3 | 50.7 | 65.5 | 56.6 | 67.9 | 72.8 | 70.0 |

Table S19: Accuracy benchmark for the test species of the Tiberius Chlorophyta model compared to *ab initio* prediction of Helixer and evidence supported prediction of BRAKER3, where BRAKER3 used the proteomes of Tiberius' training species and short-read RNA-Seq.

| Species | Tool | Exon | | | Transcript | | | Gene | | |
|---|---|---|---|---|---|---|---|---|---|---|
| | | Sn | Pr | F1 | Sn | Pr | F1 | Sn | Pr | F1 |
| *Bathycoccus prasinos* | Helixer | 52.9 | 52.4 | 52.6 | 52.7 | 59.2 | 55.8 | 52.7 | 59.2 | 55.8 |
| | Tiberius | 48.4 | 51.1 | 49.7 | 48.2 | 52.1 | 50.1 | 48.2 | 52.1 | 50.1 |
| | BRAKER3 | 41.2 | 44.4 | 42.7 | 38.4 | 41.0 | 39.7 | 38.5 | 51.2 | 44.0 |
| *Edaphochlamys debaryana* | Helixer | 86.1 | 75.9 | 80.7 | 33.8 | 27.5 | 30.3 | 33.8 | 27.5 | 30.3 |
| | Tiberius | 84.3 | 82.8 | 83.5 | 41.2 | 37.6 | 39.3 | 41.2 | 37.6 | 39.3 |
| | BRAKER3 | 61.1 | 89.3 | 72.6 | 45.6 | 52.6 | 48.9 | 45.7 | 60.1 | 51.9 |

Table S20: Accuracy benchmark for the test species of the Tiberius Bacillariophyta model compared to evidence supported prediction of BRAKER3, where BRAKER3 used the proteomes of Tiberius' training species and short-read RNA-Seq.

| Species | Tool | Exon | | | Transcript | | | Gene | | |
|---|---|---|---|---|---|---|---|---|---|---|
| | | Sn | Pr | F1 | Sn | Pr | F1 | Sn | Pr | F1 |
| *Phaeodactylum tricornutum* | Tiberius | 58.7 | 70.0 | 63.9 | 57.7 | 62.7 | 60.1 | 57.7 | 62.7 | 60.1 |
| | BRAKER3 | 59.8 | 67.3 | 63.3 | 60.3 | 58.2 | 59.2 | 60.4 | 69.5 | 64.6 |
| *Thalassiosira pseudonana* | Tiberius | 74.8 | 80.0 | 77.3 | 62.7 | 69.8 | 66.1 | 67.0 | 69.8 | 68.4 |
| | BRAKER3 | 60.3 | 77.8 | 67.9 | 54.0 | 62.1 | 57.8 | 57.6 | 76.3 | 65.6 |

Table S21: Accuracy benchmark for the test species of the Tiberius Mesangiospermae model compared to *ab initio* prediction of Helixer and ANNEVO, and evidence supported prediction of BRAKER3, where BRAKER3 used the proteomes of Tiberius' training species and short-read RNA-Seq.

| Species | Tool | Exon | | | Transcript | | | Gene | | |
|---|---|---|---|---|---|---|---|---|---|---|
| | | Sn | Pr | F1 | Sn | Pr | F1 | Sn | Pr | F1 |
| *Arabidopsis thaliana* | Helixer | 82.2 | 89.5 | 85.7 | 50.3 | 74.1 | 59.9 | 72.7 | 74.1 | 73.4 |
| | ANNEVO | 78.6 | 88.9 | 83.4 | 43.3 | 69.0 | 53.2 | 62.7 | 69.0 | 65.7 |
| | Tiberius | 82.4 | 95.8 | 88.6 | 53.7 | 85.7 | 66.0 | 77.7 | 85.7 | 81.5 |
| | BRAKER3 | 80.9 | 94.7 | 87.3 | 55.9 | 77.9 | 65.1 | 78.6 | 83.9 | 81.2 |
| *Brachypodium distachyon* | Helixer | 74.6 | 77.6 | 76.1 | 38.0 | 53.6 | 44.5 | 49.6 | 53.6 | 51.5 |
| | ANNEVO | 69.8 | 85.1 | 76.7 | 32.9 | 60.1 | 42.5 | 42.9 | 60.1 | 50.1 |
| | Tiberius | 81.3 | 88.9 | 84.9 | 54.0 | 64.3 | 58.7 | 68.0 | 64.3 | 66.1 |
| | BRAKER3 | 67.5 | 87.5 | 76.2 | 42.1 | 56.6 | 48.3 | 54.2 | 60.4 | 57.1 |
| *Eschscholzia californica* | Helixer | 82.7 | 75.1 | 78.7 | 43.1 | 52.4 | 47.3 | 61.0 | 52.4 | 56.4 |
| | ANNEVO | 78.6 | 84.8 | 81.6 | 39.0 | 61.7 | 47.8 | 55.2 | 61.7 | 58.3 |
| | Tiberius | 83.4 | 91.6 | 87.3 | 51.9 | 75.4 | 61.5 | 73.5 | 75.4 | 74.4 |
| | BRAKER3 | 79.1 | 94.6 | 86.2 | 50.9 | 79.2 | 62.0 | 70.0 | 85.6 | 77.0 |
| *Freycinetia multiflora* | Helixer | 86.5 | 67.7 | 76.0 | 47.6 | 39.6 | 43.2 | 47.6 | 39.6 | 43.2 |
| | ANNEVO | 83.1 | 77.5 | 80.2 | 49.0 | 49.0 | 49.0 | 49.0 | 49.0 | 49.0 |
| | Tiberius | 81.9 | 85.2 | 83.5 | 48.8 | 56.4 | 52.3 | 65.2 | 56.4 | 60.5 |
| | BRAKER3 | 81.5 | 92.1 | 86.5 | 67.3 | 69.6 | 68.4 | 67.3 | 79.9 | 73.1 |
| *Medicago truncatula* | Helixer | 85.7 | 73.7 | 79.2 | 54.1 | 49.9 | 51.9 | 63.8 | 49.9 | 56.0 |
| | ANNEVO | 82.9 | 83.9 | 83.4 | 50.8 | 60.8 | 55.4 | 59.9 | 60.8 | 60.3 |
| | Tiberius | 84.7 | 89.4 | 87.0 | 60.5 | 71.1 | 65.4 | 71.3 | 71.1 | 71.2 |
| | BRAKER3 | 82.1 | 92.0 | 86.8 | 64.1 | 73.5 | 68.5 | 73.0 | 79.2 | 76.0 |
| *Mimulus guttatus* | Helixer | 89.2 | 84.4 | 86.7 | 60.8 | 66.4 | 63.5 | 74.3 | 66.4 | 70.1 |
| | ANNEVO | 85.2 | 84.9 | 85.0 | 52.7 | 62.4 | 57.1 | 64.3 | 62.4 | 63.3 |
| | Tiberius | 90.3 | 91.6 | 90.9 | 68.9 | 75.9 | 72.2 | 84.2 | 75.9 | 79.8 |
| | BRAKER3 | 87.9 | 94.2 | 90.9 | 69.0 | 79.2 | 73.7 | 82.2 | 84.9 | 83.5 |
| *Urochloa brizantha* | Helixer | 87.6 | 62.2 | 72.7 | 62.0 | 43.1 | 50.9 | 62.0 | 43.1 | 50.9 |
| | ANNEVO | 82.3 | 79.3 | 80.8 | 54.4 | 56.2 | 55.3 | 54.4 | 56.2 | 55.3 |
| | Tiberius | 83.2 | 84.3 | 83.7 | 59.3 | 60.2 | 59.7 | 67.7 | 60.2 | 63.7 |
| | BRAKER3 | 79.2 | 88.9 | 83.8 | 65.7 | 66.2 | 65.9 | 65.7 | 72.2 | 68.8 |

Table S22: Accuracy benchmark for the test species of the Tiberius Fungi model compared to *ab initio* prediction of Helixer and ANNEVO, and evidence supported prediction of BRAKER3, where BRAKER3 used the proteomes of Tiberius' training species and short-read RNA-Seq.

| Species | Tool | Exon | | | Transcript | | | Gene | | |
|---|---|---|---|---|---|---|---|---|---|---|
| | | Sn | Pr | F1 | Sn | Pr | F1 | Sn | Pr | F1 |
| *Agaricus bisporus* | Helixer | 67.6 | 57.9 | 62.4 | 23.1 | 18.7 | 20.7 | 23.1 | 18.7 | 20.7 |
| | ANNEVO | 74.2 | 68.4 | 71.2 | 32.7 | 31.1 | 31.9 | 32.7 | 31.1 | 31.9 |
| | Tiberius | 79.6 | 77.5 | 78.5 | 44.0 | 44.3 | 44.1 | 44.0 | 44.3 | 44.1 |
| | BRAKER3 | 74.6 | 78.2 | 76.4 | 45.7 | 45.4 | 45.5 | 45.7 | 52.4 | 48.8 |
| *Cryphonectria parasitica* | Helixer | 59.9 | 52.3 | 55.8 | 39.7 | 32.1 | 35.5 | 39.7 | 32.1 | 35.5 |
| | ANNEVO | 59.3 | 58.1 | 58.7 | 39.8 | 36.3 | 38.0 | 39.8 | 36.3 | 38.0 |
| | Tiberius | 62.3 | 65.6 | 63.9 | 45.2 | 43.6 | 44.4 | 45.2 | 43.6 | 44.4 |
| | BRAKER3 | 59.8 | 65.6 | 62.6 | 45.5 | 41.9 | 43.6 | 45.5 | 47.9 | 46.7 |
| *Parastagonospora nodorum* | Helixer | 37.5 | 38.0 | 37.7 | 22.3 | 24.2 | 23.2 | 22.3 | 24.2 | 23.2 |
| | ANNEVO | 43.0 | 47.0 | 44.9 | 27.9 | 32.5 | 30.0 | 27.9 | 32.5 | 30.0 |
| | Tiberius | 45.6 | 53.7 | 49.3 | 32.0 | 38.8 | 35.1 | 32.0 | 38.8 | 35.1 |
| | BRAKER3 | 38.6 | 54.3 | 45.1 | 29.4 | 38.3 | 33.3 | 29.4 | 42.4 | 34.7 |
| *Puccinia striiformis* | Helixer | 66.1 | 43.2 | 52.3 | 28.5 | 19.4 | 23.1 | 28.5 | 19.4 | 23.1 |
| | ANNEVO | 58.6 | 54.0 | 56.2 | 24.9 | 24.2 | 24.5 | 24.9 | 24.2 | 24.5 |
| | Tiberius | 67.9 | 66.3 | 67.1 | 38.4 | 37.3 | 37.8 | 38.4 | 37.3 | 37.8 |
| | BRAKER3 | 64.5 | 85.1 | 73.4 | 47.9 | 59.3 | 53.0 | 48.0 | 66.7 | 55.8 |
| *Punctularia strigosozonata* | Helixer | 74.7 | 61.1 | 67.2 | 30.8 | 24.6 | 27.4 | 30.8 | 24.6 | 27.4 |
| | ANNEVO | 76.9 | 68.2 | 72.3 | 35.0 | 31.5 | 33.2 | 35.0 | 31.5 | 33.2 |
| | Tiberius | 82.3 | 77.5 | 79.8 | 47.3 | 46.0 | 46.6 | 47.4 | 46.0 | 46.7 |
| | BRAKER3 | 72.4 | 78.5 | 75.3 | 45.5 | 46.3 | 45.9 | 45.5 | 52.2 | 48.6 |
| *Tilletiopsis washingtonensis* | Helixer | 83.4 | 70.6 | 76.5 | 48.9 | 40.6 | 44.4 | 48.9 | 40.6 | 44.4 |
| | ANNEVO | 84.7 | 80.8 | 82.7 | 53.0 | 56.2 | 54.6 | 53.0 | 56.2 | 54.6 |
| | Tiberius | 87.9 | 85.0 | 86.4 | 61.3 | 66.4 | 63.7 | 61.3 | 66.4 | 63.7 |
| | BRAKER3 | 72.8 | 85.3 | 78.6 | 48.5 | 59.7 | 53.5 | 48.5 | 66.6 | 56.1 |

Table S23: Accuracy benchmark for the test species of the Tiberius Insecta model compared to *ab initio* prediction of Helixer and ANNEVO, and evidence supported prediction of BRAKER3, where BRAKER3 used the proteomes of Tiberius' training species and short-read RNA-Seq.

| Species | Tool | Exon | | | Transcript | | | Gene | | |
|---|---|---|---|---|---|---|---|---|---|---|
| | | Sn | Pr | F1 | Sn | Pr | F1 | Sn | Pr | F1 |
| *Bombyx mori* | Helixer | 80.1 | 73.0 | 76.4 | 20.3 | 30.9 | 24.5 | 34.7 | 30.9 | 32.7 |
| | ANNEVO | 80.2 | 80.0 | 80.1 | 24.2 | 40.5 | 30.3 | 41.2 | 40.5 | 40.8 |
| | Tiberius | 77.8 | 85.6 | 81.5 | 35.4 | 41.9 | 38.4 | 60.2 | 41.9 | 49.4 |
| | BRAKER3 | 81.7 | 94.2 | 87.5 | 48.7 | 71.7 | 58.0 | 73.1 | 79.7 | 76.3 |
| *Cataglyphis hispanica* | Helixer | 81.5 | 80.9 | 81.2 | 31.0 | 45.4 | 36.8 | 51.8 | 45.4 | 48.4 |
| | ANNEVO | 80.8 | 83.5 | 82.1 | 31.2 | 50.5 | 38.6 | 52.1 | 50.5 | 51.3 |
| | Tiberius | 84.1 | 86.0 | 85.0 | 43.3 | 47.8 | 45.4 | 72.4 | 47.8 | 57.6 |
| | BRAKER3 | 85.4 | 88.8 | 87.1 | 57.9 | 56.1 | 57.0 | 81.1 | 60.3 | 69.2 |
| *Colias croceus* | Helixer | 85.0 | 75.1 | 79.7 | 30.5 | 35.5 | 32.8 | 42.6 | 35.5 | 38.7 |
| | ANNEVO | 83.6 | 82.9 | 83.2 | 33.8 | 47.3 | 39.4 | 47.2 | 47.3 | 47.2 |
| | Tiberius | 87.2 | 90.6 | 88.9 | 49.6 | 60.3 | 54.4 | 69.3 | 60.3 | 64.5 |
| | BRAKER3 | 82.8 | 93.2 | 87.7 | 59.4 | 67.9 | 63.4 | 73.8 | 77.6 | 75.7 |
| *Danaus plexippus* | Helixer | 84.8 | 79.5 | 82.1 | 32.4 | 41.9 | 36.5 | 48.5 | 41.9 | 45.0 |
| | ANNEVO | 83.7 | 83.9 | 83.8 | 34.3 | 50.7 | 40.9 | 51.4 | 50.7 | 51.0 |
| | Tiberius | 87.5 | 91.5 | 89.5 | 49.4 | 62.1 | 55.0 | 73.9 | 62.1 | 67.5 |
| | BRAKER3 | 86.7 | 93.5 | 90.0 | 61.8 | 66.9 | 64.2 | 80.8 | 74.4 | 77.5 |
| *Drosophila melanogaster* | Helixer | 79.2 | 84.1 | 81.6 | 43.1 | 66.0 | 52.1 | 68.9 | 66.0 | 67.4 |
| | ANNEVO | 77.5 | 86.0 | 81.5 | 41.8 | 68.5 | 51.9 | 66.7 | 68.5 | 67.6 |
| | Tiberius | 81.1 | 90.7 | 85.6 | 49.8 | 69.5 | 58.0 | 79.6 | 69.5 | 74.2 |
| | BRAKER3 | 83.8 | 94.9 | 89.0 | 60.4 | 80.6 | 69.1 | 85.7 | 89.6 | 87.6 |
| *Leptidea sinapis* | Helixer | 78.9 | 59.6 | 67.9 | 18.3 | 21.4 | 19.7 | 28.9 | 21.4 | 24.6 |
| | ANNEVO | 80.6 | 70.2 | 75.0 | 25.5 | 33.8 | 29.1 | 40.3 | 33.8 | 36.8 |
| | Tiberius | 80.3 | 74.3 | 77.2 | 38.2 | 30.9 | 34.2 | 60.1 | 30.9 | 40.8 |
| | BRAKER3 | 75.1 | 86.5 | 80.4 | 48.0 | 61.8 | 54.0 | 68.5 | 68.0 | 68.2 |
| *Nymphalis io* | Helixer | 84.2 | 74.3 | 78.9 | 27.8 | 35.5 | 31.2 | 42.9 | 35.5 | 38.9 |
| | ANNEVO | 84.5 | 80.3 | 82.3 | 33.7 | 46.6 | 39.1 | 52.1 | 46.6 | 49.2 |
| | Tiberius | 87.2 | 86.9 | 87.0 | 46.2 | 50.1 | 48.1 | 71.3 | 50.1 | 58.8 |
| | BRAKER3 | 85.2 | 94.5 | 89.6 | 59.3 | 71.9 | 65.0 | 79.9 | 81.3 | 80.6 |
| *Osmia bicornis* | Helixer | 77.2 | 77.7 | 77.4 | 26.4 | 41.7 | 32.3 | 43.9 | 41.7 | 42.8 |
| | ANNEVO | 75.8 | 81.2 | 78.4 | 26.7 | 47.1 | 34.1 | 44.4 | 47.1 | 45.7 |
| | Tiberius | 80.6 | 79.2 | 79.9 | 40.2 | 36.4 | 38.2 | 66.8 | 36.4 | 47.1 |
| | BRAKER3 | 81.2 | 89.7 | 85.2 | 51.3 | 58.1 | 54.5 | 72.5 | 64.6 | 68.3 |
| *Tribolium castaneum* | Helixer | 53.3 | 48.1 | 50.6 | 13.2 | 15.7 | 14.3 | 20.9 | 15.7 | 17.9 |
| | ANNEVO | 68.5 | 74.2 | 71.2 | 25.2 | 39.0 | 30.6 | 39.8 | 39.0 | 39.4 |
| | Tiberius | 74.6 | 81.8 | 78.0 | 38.3 | 46.7 | 42.1 | 60.4 | 46.7 | 52.7 |
| | BRAKER3 | 83.2 | 90.7 | 86.8 | 57.7 | 67.4 | 62.2 | 80.2 | 74.6 | 77.3 |
| *Vanessa cardui* | Helixer | 85.2 | 75.0 | 79.8 | 31.9 | 37.8 | 34.6 | 43.7 | 37.8 | 40.5 |
| | ANNEVO | 85.4 | 84.1 | 84.7 | 38.4 | 51.8 | 44.1 | 52.5 | 51.8 | 52.1 |
| | Tiberius | 88.0 | 87.1 | 87.5 | 50.8 | 50.5 | 50.6 | 69.5 | 50.5 | 58.5 |
| | BRAKER3 | 84.0 | 93.8 | 88.6 | 60.3 | 68.9 | 64.3 | 73.6 | 78.3 | 75.9 |
| *Zerene cesonia* | Helixer | 85.1 | 73.9 | 79.1 | 30.4 | 34.1 | 32.1 | 41.0 | 34.1 | 37.2 |
| | ANNEVO | 84.5 | 80.8 | 82.6 | 35.2 | 44.7 | 39.4 | 47.6 | 44.7 | 46.1 |
| | Tiberius | 87.3 | 89.3 | 88.3 | 48.9 | 59.0 | 53.5 | 66.0 | 59.0 | 62.3 |
| | BRAKER3 | 73.8 | 93.1 | 82.3 | 59.6 | 66.6 | 62.9 | 71.5 | 73.7 | 72.6 |

Table S24: Accuracy benchmark for the test species of the Tiberius Mammalia model compared to *ab initio* prediction of Helixer and ANNEVO, and evidence supported prediction of BRAKER3, where BRAKER3 used the proteomes of Tiberius' training species and short-read RNA-Seq.

| Species | Tool | Exon | | | Transcript | | | Gene | | |
|---|---|---|---|---|---|---|---|---|---|---|
| | | Sn | Pr | F1 | Sn | Pr | F1 | Sn | Pr | F1 |
| *Bos taurus* | Helixer | 76.2 | 70.7 | 73.3 | 10.2 | 24.0 | 14.3 | 23.1 | 24.0 | 23.5 |
| | ANNEVO | 81.3 | 82.8 | 82.0 | 20.3 | 41.4 | 27.2 | 46.2 | 41.4 | 43.7 |
| | Tiberius | 82.5 | 90.2 | 86.2 | 27.3 | 57.8 | 37.1 | 62.2 | 57.8 | 59.9 |
| | BRAKER3 | 67.2 | 94.9 | 78.7 | 32.9 | 71.3 | 45.0 | 64.4 | 77.0 | 70.1 |
| *Delphinapterus leucas* | Helixer | 77.3 | 72.5 | 74.8 | 10.6 | 23.0 | 14.5 | 23.3 | 23.0 | 23.1 |
| | ANNEVO | 80.5 | 77.5 | 79.0 | 17.8 | 30.0 | 22.3 | 39.3 | 30.0 | 34.0 |
| | Tiberius | 82.6 | 91.8 | 87.0 | 27.3 | 59.4 | 37.4 | 60.3 | 59.4 | 59.8 |
| | BRAKER3 | 66.6 | 94.1 | 78.0 | 34.6 | 65.8 | 45.4 | 65.8 | 72.4 | 68.9 |
| *Homo sapiens* | Helixer | 78.8 | 69.8 | 74.0 | 10.5 | 25.0 | 14.8 | 27.4 | 25.0 | 26.1 |
| | ANNEVO | 82.8 | 80.5 | 81.6 | 18.3 | 38.4 | 24.8 | 47.7 | 38.4 | 42.5 |
| | Tiberius | 85.0 | 90.2 | 87.5 | 27.4 | 61.2 | 37.9 | 71.4 | 61.2 | 65.9 |
| | BRAKER3 | 74.0 | 92.0 | 82.0 | 34.3 | 61.5 | 44.0 | 76.5 | 68.4 | 72.2 |

Table S25: Accuracy benchmark for the non-mammalian test species of the Tiberius Vertebrata model compared to *ab initio* prediction of Helixer and ANNEVO, and evidence supported prediction of BRAKER3, where BRAKER3 used the proteomes of Tiberius' training species and short-read RNA-Seq.

| Species | Tool | Exon | | | Transcript | | | Gene | | |
|---|---|---|---|---|---|---|---|---|---|---|
| | | Sn | Pr | F1 | Sn | Pr | F1 | Sn | Pr | F1 |
| *Archocentrus centrarchus* | Helixer | 82.2 | 68.7 | 74.8 | 16.9 | 20.7 | 18.6 | 24.2 | 20.7 | 22.3 |
| | ANNEVO | 84.1 | 80.7 | 82.4 | 30.0 | 40.6 | 34.5 | 43.0 | 40.6 | 41.8 |
| | Tiberius | 88.2 | 86.1 | 87.1 | 44.9 | 53.6 | 48.9 | 64.3 | 53.6 | 58.5 |
| | BRAKER3 | 76.9 | 94.3 | 84.7 | 55.4 | 69.7 | 61.7 | 69.4 | 79.4 | 74.1 |
| *Betta splendens* | Helixer | 82.5 | 79.0 | 80.7 | 16.9 | 30.0 | 21.6 | 32.3 | 30.0 | 31.1 |
| | ANNEVO | 84.2 | 86.5 | 85.3 | 24.9 | 47.0 | 32.6 | 47.6 | 47.0 | 47.3 |
| | Tiberius | 88.8 | 93.5 | 91.1 | 37.8 | 68.3 | 48.7 | 72.2 | 68.3 | 70.2 |
| | BRAKER3 | 84.5 | 96.4 | 90.1 | 48.3 | 73.8 | 58.4 | 77.9 | 84.6 | 81.1 |
| *Gallus gallus* | Helixer | 78.1 | 72.2 | 75.0 | 10.2 | 25.7 | 14.6 | 26.3 | 25.7 | 26.0 |
| | ANNEVO | 77.3 | 84.5 | 80.7 | 14.6 | 40.3 | 21.4 | 37.6 | 40.3 | 38.9 |
| | Tiberius | 82.1 | 89.9 | 85.8 | 23.4 | 57.9 | 33.3 | 60.3 | 57.9 | 59.1 |
| | BRAKER3 | 76.7 | 96.2 | 85.4 | 32.9 | 73.6 | 45.5 | 69.9 | 82.7 | 75.8 |
| *Pristiophorus japonicus* | Helixer | 67.6 | 47.0 | 55.4 | 5.8 | 6.4 | 6.1 | 9.6 | 6.4 | 7.7 |
| | ANNEVO | 77.9 | 66.2 | 71.6 | 15.3 | 17.5 | 16.3 | 25.4 | 17.5 | 20.7 |
| | Tiberius | 80.3 | 67.5 | 73.3 | 23.5 | 17.0 | 19.7 | 38.9 | 17.0 | 23.7 |
| | BRAKER3 | 62.3 | 94.3 | 75.0 | 36.8 | 65.4 | 47.1 | 52.7 | 73.3 | 61.3 |
| *Takifugu rubripes* | Helixer | 80.9 | 75.7 | 78.2 | 15.1 | 24.8 | 18.8 | 27.2 | 24.8 | 25.9 |
| | ANNEVO | 82.9 | 83.4 | 83.1 | 23.7 | 41.4 | 30.1 | 42.7 | 41.4 | 42.0 |
| | Tiberius | 87.1 | 91.2 | 89.1 | 37.0 | 61.8 | 46.3 | 66.6 | 61.8 | 64.1 |
| | BRAKER3 | 80.3 | 96.2 | 87.5 | 48.3 | 72.7 | 58.0 | 75.1 | 83.8 | 79.2 |
| *Zootoca vivipara* | Helixer | 81.2 | 66.4 | 73.1 | 11.8 | 18.6 | 14.4 | 20.3 | 18.6 | 19.4 |
| | ANNEVO | 83.4 | 78.8 | 81.0 | 21.4 | 33.0 | 26.0 | 36.9 | 33.0 | 34.8 |
| | Tiberius | 87.3 | 89.0 | 88.1 | 34.8 | 52.5 | 41.9 | 60.0 | 52.5 | 56.0 |
| | BRAKER3 | 80.3 | 93.8 | 86.5 | 49.1 | 65.7 | 56.2 | 71.7 | 77.0 | 74.3 |

Table S26: Accuracy comparison between the Tiberius version v1.1.8 and v2.0.0 with reimplemented core. The table shows the average accuracy across all test species for each Tiberius model.

| Clade | Tool | $n$ | Exon | | | Transcript | | | Gene | | |
|---|---|---|---|---|---|---|---|---|---|---|---|
| | | | Sn | Pr | F1 | Sn | Pr | F1 | Sn | Pr | F1 |
| Chlorophyta | Tiberius v1.1.8 | 2 | 66.2 | 67.0 | 66.6 | 44.6 | 44.9 | 44.7 | 44.6 | 44.9 | 44.7 |
| | Tiberius v2.0.0 | 2 | 66.3 | 67.0 | 66.6 | 44.7 | 44.9 | 44.7 | 44.7 | 44.9 | 44.7 |
| Bacillariophyta | Tiberius v1.1.8 | 2 | 66.7 | 75.1 | 70.6 | 60.2 | 66.3 | 63.1 | 62.2 | 66.3 | 64.2 |
| | Tiberius v2.0.0 | 2 | 66.8 | 75.0 | 70.6 | 60.2 | 66.2 | 63.1 | 62.4 | 66.2 | 64.2 |
| Mesangiospermae | Tiberius v1.1.8 | 7 | 82.1 | 90.8 | 86.1 | 56.6 | 71.3 | 62.3 | 66.5 | 71.3 | 68.6 |
| | Tiberius v2.0.0 | 7 | 82.1 | 90.8 | 86.1 | 56.6 | 71.2 | 62.2 | 66.5 | 71.2 | 68.6 |
| Fungi | Tiberius v1.1.8 | 6 | 70.8 | 71.0 | 70.8 | 44.7 | 46.2 | 45.3 | 44.7 | 46.2 | 45.3 |
| | Tiberius v2.0.0 | 6 | 70.9 | 70.9 | 70.9 | 44.7 | 46.1 | 45.3 | 44.7 | 46.1 | 45.3 |
| Insecta | Tiberius v1.1.8 | 11 | 83.5 | 86.1 | 84.7 | 44.9 | 51.6 | 47.8 | 68.7 | 51.6 | 58.5 |
| | Tiberius v2.0.0 | 11 | 83.2 | 85.7 | 84.4 | 44.6 | 50.5 | 47.1 | 68.1 | 50.5 | 57.6 |
| Mammalia | Tiberius v1.1.8 | 3 | 82.4 | 90.8 | 86.4 | 27.3 | 59.7 | 37.4 | 64.5 | 59.7 | 62.0 |
| | Tiberius v2.0.0 | 3 | 83.4 | 90.7 | 86.9 | 27.3 | 59.5 | 37.4 | 64.6 | 59.5 | 61.9 |
| Vertebrata | Tiberius v1.1.8 | 6 | 85.3 | 86.2 | 85.6 | 33.5 | 51.9 | 39.8 | 60.3 | 51.9 | 55.3 |
| | Tiberius v2.0.0 | 6 | 85.6 | 86.2 | 85.8 | 33.6 | 51.9 | 39.8 | 60.4 | 51.9 | 55.3 |
| Overall | Tiberius v1.1.8 | 37 | 79.5 | 83.3 | 81.3 | 44.6 | 55.6 | 48.6 | 61.0 | 55.6 | 57.6 |
| | Tiberius v2.0.0 | 37 | 79.6 | 83.2 | 81.2 | 44.5 | 55.2 | 48.4 | 60.9 | 55.2 | 57.3 |

Table S27: Runtime comparison between the Tiberius v1.1.8 and v2.0.0 with reimplemented core.

| Clade | Tool | $n$ | Runtime (h:mm) |
|---|---|---|---|
| Chlorophyta | Tiberius v.1.1.8 | 2 | 0:04 |
| | Tiberius v2.0.0 | 2 | 0:03 |
| Bacillariophyta | Tiberius v.1.1.8 | 2 | 0:02 |
| | Tiberius v2.0.0 | 2 | 0:01 |
| Mesangiospermae | Tiberius v.1.1.8 | 7 | 0:32 |
| | Tiberius v2.0.0 | 7 | 0:21 |
| Fungi | Tiberius v.1.1.8 | 6 | 0:02 |
| | Tiberius v2.0.0 | 6 | 0:02 |
| Insecta | Tiberius v.1.1.8 | 11 | 0:18 |
| | Tiberius v2.0.0 | 11 | 0:12 |
| Mammalia | Tiberius v.1.1.8 | 3 | 1:58 |
| | Tiberius v2.0.0 | 3 | 1:20 |
| Vertebrata | Tiberius v.1.1.8 | 6 | 1:22 |
| | Tiberius v2.0.0 | 6 | 0:58 |
| Overall | Tiberius v.1.1.8 | 37 | 0:35 |
| | Tiberius v2.0.0 | 37 | 0:24 |

Table S28: The names of the models used for Tiberius, ANNEVO and Helixer for the benchmark on the test species for each clade.

| Clade | Tiberius model | ANNEVO model | Helixer model |
|---|---|---|---|
| Chlorophyta | chlorophyta | | land_plant |
| Bacillariophyta | diatoms_unmasked | | |
| Mesangiospermae | angiosperms | ANNEVO_Embryophyta.pt | land_plant |
| Fungi | fungi | ANNEVO_Fungi.pt | fungi |
| Insecta | insecta_unmasked_v2 | ANNEVO_Invertebrate.pt | invertebrate |
| Mammalia | mammalia_nosofttmasking_v2 | ANNEVO_Mammalia2.pt | vertebrate |
| Vertebrata | vertebrate | ANNEVO_Vertebrate_other.pt | vertebrate |

| Model / Clade | GPU | Phase 2 (HMM training) |
|---|---|---|
| Chlorophyta | A100 (80 GB) | Yes |
| Bacillariophyta | A100 (80 GB) | Yes |
| Mesangiospermae | A100 (40 GB) | No |
| Insecta | A100 (80 GB) | Yes |
| Fungi | A100 (80 GB) | Yes |
| Mammalia | A100 (80 GB) | Yes |
| Vertebrata | 4x NVIDIA H100 (64 GB VRAM each) | Yes |

Table S29: Overview of the GPU hardware used for training each model and whether training included a second phase with the HMM layer.

| Model / Clade | CNN units | LSTM units | Params (M) |
|---|---|---|---|
| Chlorophyta | 128 | 744 | 8.0 |
| Bacillariophyta | 128 | 744 | 8.0 |
| Mesangiospermae | 128 | 744 | 8.0 |
| Insecta | 128 | 372 | 2.5 |
| Fungi | 128 | 744 | 8.0 |
| Vertebrata | 256 | 744 | 9.8 |

Table S30: Model sizes for all trained clade-specific Tiberius models. The overall architecture is identical to the original mammalian Tiberius model, consisting of three CNN followed by two biLSTM layers. Only layer sizes (CNN units/channels and LSTM units) were varied between the models, resulting in different total parameter counts.

| Level | Comparator | $\overline{F1}_{\text{Tib}}$ | $\overline{F1}_{\text{comp}}$ | median $\Delta$ | wins:losses:ties | $p$ |
|---|---|---|---|---|---|---|
| Exon | Helixer | 82.9 | 72.8 | +9.3 | 33:0:0 | $2.33 \times 10^{-10}$ |
| Exon | ANNEVO | 82.9 | 77.8 | +5.2 | 33:0:0 | $2.33 \times 10^{-10}$ |
| Transcript | Helixer | 47.8 | 30.9 | +16.0 | 33:0:0 | $2.33 \times 10^{-10}$ |
| Transcript | ANNEVO | 47.8 | 37.1 | +11.9 | 33:0:0 | $2.33 \times 10^{-10}$ |
| Gene | Helixer | 58.2 | 36.8 | +18.0 | 33:0:0 | $2.33 \times 10^{-10}$ |
| Gene | ANNEVO | 58.2 | 45.1 | +13.3 | 33:0:0 | $2.33 \times 10^{-10}$ |

Table S31: Two-sided paired Wilcoxon signed-rank tests comparing Tiberius against Helixer and ANNEVO on the per-genome F1 score across the $n = 33$ benchmark genomes, at exon, transcript and gene level. Columns show the mean F1 of Tiberius ($\overline{F1}_{\text{Tib}}$) and of the comparator ($\overline{F1}_{\text{comp}}$), the median paired difference ($\Delta = F1_{\text{Tib}} - F1_{\text{comp}}$) and the number of genomes on which Tiberius has higher, lower or the same F1 score as the comparator. Because Tiberius has the higher F1 on all 33 genomes in every comparison (33:0:0), the two-sided $p$-value reaches the exact signed-rank minimum $2/2^{33} = 2.33 \times 10^{-10}$ in all cases. All six tests across the three levels and two comparators reject the null hypothesis of no per-genome difference at $p < 2.33 \times 10^{-10}$ and the sign of the median difference is positive in every case.

| Clade | Test species | Closest shared rank | LCA (NCBI taxon) | Divergence (Mya) | Closest training species at this rank |
|---|---|---|---|---|---|
| Mammalia | *B. taurus* | order | Artiodactyla (order) | 64.0 | *C. bactrianus* |
| | *D. leucas* | order | Artiodactyla (order) | 64.0 | *C. bactrianus* |
| | *H. sapiens* | order | Simiiformes (infraorder) | 42.9 | *A. nancymaae*, *C. capucinus*, *S. boliviensis* |
| Vertebrata (non-mammalian) | *A. centrarchus* | class | Ovalentaria (clade) | 87.2 | *P. ranga* |
| | *B. splendens* | class | Percomorphaceae (clade) | 112.5 | *L. mixtus*, *P. ranga*, *S. solea* (+1 more) |
| | *G. gallus* | class | Neognathae (infraclass) | 90.8 | *A. apus*, *C. hawaiiensis*, *F. peregrinus* (+1 more) |
| | *P. japonicus* | class | Selachii (infraclass) | 225.0 | *C. carcharias* |
| | *T. rubripes* | class | Eupercaria (clade) | 99.6 | *L. mixtus*, *S. aurata* |
| | *Z. vivipara* | family | Lacertinae (subfamily) | 46.0 | *P. raffonei* |
| Insecta | *B. mori* | order | Obtectomera (clade) | 95.4 | *A. ipsilon*, *A. tragopoginis*, *A. lueneburgensis* (+16 more) |
| | *C. hispanica* | family | Formicini (tribe) | 55.5 | *F. exsecta* |
| | *C. croceus* | order | Papilionoidea (superfamily) | 101.2 | *B. anynana*, *L. megera*, *M. cinxia* |
| | *D. plexippus* | family | Nymphalidae (family) | 72.9 | *B. anynana*, *L. megera*, *M. cinxia* |
| | *D. melanogaster* | **genus** | melanogaster group (species group) | 28.1 | *D. birchii*, *D. suzukii* |
| | *L. sinapis* | order | Papilionoidea (superfamily) | 101.2 | *B. anynana*, *L. megera*, *M. cinxia* |
| | *N. io* | family | Nymphalinae (subfamily) | 42.5 | *M. cinxia* |
| | *O. bicornis* | order | Anthophila (clade) | 110.3 | *A. minutula*, *A. mellifera*, *B. impatiens* (+1 more) |
| | *T. castaneum* | order | Tenebrionoidea (superfamily) | 115.0 | *H. cichorii* |
| | *V. cardui* | family | Nymphalinae (subfamily) | 42.5 | *M. cinxia* |
| | *Z. cesonia* | order | Papilionoidea (superfamily) | 101.2 | *B. anynana*, *L. megera*, *M. cinxia* |
| Mesangiospermae | *A. thaliana* | family | Brassicaceae (family) | 25.9 | *B. rapa* |
| | *B. distachyon* | family | Pooideae (subfamily) | 33.0 | *T. intermedium* |
| | *E. californica* | order | Ranunculales (order) | 107.9 | *A. coerulea* |
| | *F. multiflora* | class | Petrosaviidae (subclass) | 119.0 | *A. tequilana*, *A. donax*, *B. micrantha* (+16 more) |
| | *M. truncatula* | family | NPAAA clade (clade) | 54.6 | *G. max* |
| | *M. guttatus* | order | Lamiales (order) | 40.9 | *L. philippensis*, *M. longifolia* |
| | *U. brizantha* | family | Paniceae (tribe) | 14.3 | *P. virgatum*, *S. viridis* |
| Fungi | *A. bisporus* | order | Agaricineae (suborder) | n/a | *P. cubensis* |
| | *C. parasitica* | order | Diaporthales (order) | n/a | *D. amygdali* |
| | *P. nodorum* | order | Pleosporineae (suborder) | 115.8 | *A. atra*, *A. burnsii*, *A. rosae* (+9 more) |
| | *P. striiformis* | phylum | Pucciniomycotina (subphylum) | n/a | *C. minutum*, *R. toruloides* |
| | *P. strigosozonata* | class | Agaricomycetes incertae sedis (no rank) | 145.3 | *D. squalens*, *F. mediterranea*, *G. trabeum* (+6 more) |
| | *T. washingtonensis* | phylum | Ustilaginomycotina (subphylum) | 280.6 | *P. flocculosa*, *P. hubeiensis*, *U. hordei* (+1 more) |
| Chlorophyta | *E. debaryana* | order | Chlamydomonadales (order) | 227.0 | *A. gubernaculifera*, *C. reinhardtii*, *D. primolecta* (+1 more) |
| | *B. prasinos* | family | Bathycoccaceae (family) | n/a | *O. lucimarinus* |
| Bacillariophyta | *T. pseudonana* | **genus** | Thalassiosira (genus) | 42.0 | *T. mediterranea* |
| | *P. tricornutum* | class | Bacillariophycidae (clade) | n/a | *C. closterium*, *M. pseudoterrestris*, *P. multistriata* (+1 more) |

Table S32: Taxonomic distance between each test species and the closest training species per clade, using NCBI Taxonomy and TimeTree [7]. For every test species we report the deepest taxonomic rank at which it shares a last common ancestor (LCA) with any training species (*Closest shared rank*), the corresponding LCA taxon, the divergence time in Mya taken as the median across all TimeTree study estimates for that species pair, and the closest training species (*Closest training species at this rank*). If more than three such training species exist, three are listed with the remainder counted. Ranks in bold indicate a shared genus or lower; no test species shares a species with any training species. Only *Thalassiosira pseudonana* (Bacillariophyta) and *Drosophila melanogaster* (Insecta) share a genus with a training species; the corresponding TimeTree divergences are 42 and 28 Mya, respectively. Five test species (mostly microbial eukaryotes and Fungi) have no pairwise studies in TimeTree and are shown as "n/a".

### References


[1] The UniProt Consortium. UniProt: the universal protein knowledgebase in 2025. *Nucleic Acids Research*, 53(D1):D609–D617, 2025.

[2] Benjamin Buchfink, Klaus Reuter, and Hajk-Georg Drost. Sensitive protein alignments at tree-of-life scale using DIAMOND. *Nature Methods*, 18(4):366–368, 2021.

[3] Gina V Filloramo, Bruce A Curtis, Emma Blanche, and John M Archibald. Re-examination of two diatom reference genomes using long-read sequencing. *BMC Genomics*, 22(1):379, 2021.

[4] Natalia Nenasheva, Clara Pitzschel, Cynthia N Webster, Alexander J Hart, Jill L Wegrzyn, Mia M Bengtsson, and Katharina J Hoff. Annotation of protein-coding genes in 49 diatom genomes from the Bacillariophyta clade. *Scientific Data*, 12(1):985, 2025.

[5] Rory J. Craig, Ahmed R. Hasan, Rob W. Ness, and Peter D. Keightley. Comparative genomics of *Chlamydomonas*. *The Plant Cell*, 33(4):1016–1041, 2021.

[6] Lars Gabriel, Felix Becker, Katharina J Hoff, and Mario Stanke. Tiberius: End-to-end deep learning with an HMM for gene prediction. *Bioinformatics*, 40(12):btae685, 2024.

[7] Sudhir Kumar, Michael Suleski, Jack M Craig, Adrienne E Kasprowicz, Maxwell Sanderford, Michael Li, Glen Stecher, and S Blair Hedges. TimeTree 5: An expanded resource for species divergence times. *Molecular Biology and Evolution*, 39(8):msac174, 2022.
